# Direct Cortical Inputs to Hippocampal Area CA1 Transmit Complementary Signals for Goal-directed Navigation

**DOI:** 10.1101/2022.11.10.516009

**Authors:** John C Bowler, Attila Losonczy

## Abstract

The entorhinal cortex (EC) is central to the brain’s navigation system. Its subregions are conventionally thought to compute dichotomous representations for spatial processing: medial entorhinal cortex (MEC) provides a global spatial map, while lateral entorhinal cortex (LEC) encodes specific sensory details of experience. While local recordings of EC circuits have amassed a vast catalogue of specialized cell types that could support navigation computations in the brain, we have little direct evidence for how these signals are actually transmitted outside of the EC to its primary downstream reader, the hippocampus, which itself is critical for the formation of spatial and episodic memories. Here we exploit *in vivo* sub-cellular imaging to directly record from EC axon terminals as they locally innervate hippocampal area CA1, while mice performed navigational and spatial learning tasks in virtual reality. We find both distinct and overlapping representations of task, location, and context in both MEC and LEC axons. While MEC transmitted a highly location- and context-specific code, LEC inputs were strongly biased by ongoing navigational goals and reward. Surprisingly, the position of the animal could be accurately decoded from either entorhinal subregion. Our results challenge prevailing dogma on the routing of spatial and non-spatial information from the cortex to the hippocampus, indicating that cortical interactions upstream of the hippocampus are critical for combining these processing streams to support navigation and memory.

## Introduction

The mammalian hippocampal formation plays a critical role in both spatial navigation and episodic memory (Scoville & Milner 1957, O’Keefe & Dostrovsky 1971). The most prominent hippocampal neural correlate of these phenomena is place coding, in which individual excitatory principal neurons in the hippocampus are specifically active in restricted regions of the environment, termed “place cells”, with the preferred locations referred to as a cell’s “place fields” (O’Keefe & Dostrovsky 1971). At the population level, place cells produce a map of external space (O’Keefe & Nadel 1978, Wilson & McNaughton 1993, Whittington et al. 2022). Importantly, place maps are altered by proximal cues, which can serve as local landmarks to support allothetic navigation (Deshmukh & Knierim 2013, Geiller et al. 2017); place maps have also been repeatedly demonstrated to incorporate goal-related information (Dupret et al. 2010, Zaremba et al. 2017, Gauthier & Tank 2018). Increasingly, studies have found evidence that adjacent cortical regions play important roles in shaping the spatial and mnemonic activity in the hippocampus (Burwell & Agster 2008, Nilssen et al. 2019). Indeed, the entorhinal cortex (EC), the primary cortical input region of the hippocampus, has been shown to have a diverse array of functionally-defined cell types capable of providing streams of both spatial and sensory information to the hippocampus (Knierim et al. 2014, Moser et al. 2014, Sugar & Moser 2019, Keene et al. 2016). Critically, both EC and hippocampal spatial maps exhibit experience-dependent changes even in familiar environments (Colgin et al. 2008, Kitamura et al. 2015) and also respond to salience and novelty (Cohen et al. 2017, Larkin et al. 2014, Barry et al. 2012). Recent models for place field formation in hippocampal subregion CA1, describe how highly tuned spatial input arriving from CA3 (Dong et al. 2021, Dupret et al. 2010) might be combined conjunctively with an external depolarizing event to create or refine spatial receptive fields (Bittner et al. 2015, 2017, Milstein et al. 2021, Priestley et al. 2022, Rolotti et al. 2022). The prominent direct (temporoammonic) projection from superficial, primarily layer III, EC to CA1 (Steward & Scoville 1976) (c.f. Kitamura et al. 2014) is ideally suited to provide instructive signals for place field formation and transformation in CA1. Given the malleable nature of hippocampal representations, any truly mechanistic model of the activity of principal cells in CA1 must rely on a comprehensive characterization of the activity in the EC-to-CA1 projections, as well as an understanding of how this activity changes with navigational context and task goals.

Direct recordings from neurons in superficial layers of the EC have provided some clues as to the types of activity that might participate in their CA1 projection. Traditionally, a dissociation between the medial (MEC) and lateral (LEC) subregions of the EC, has been identified. MEC neurons display range of spatial response profiles: grid cells tessellate space with hexagonally repeating receptive fields (Hafting et al. 2005), border cells are active along geometric borders (Solstad et al. 2008), and head-direction cells have bearing-dependent tuning (Sargolini et al. 2006). Additionally, speed cells modulate responses proportionally to an animal’s current velocity and are have been suggested to play a role in navigation based on path integration (Kropff et al. 2015). Intriguingly, there is evidence for more complex representations in MEC, with cells responding to conjunctions of parameters (Sargolini et al. 2006, Hardcastle et al. 2017), as well as associative schemes such as object-vector coding (Høydal et al. 2019). The high proportion of spatially modulated MEC cells suggest that MEC may be involved in representing spatial information related to global frames of reference. In contrast, LEC has been proposed to primarily process information related to individual items, such as sensory cues and locations based on a local frame of reference (Knierim et al. 2014, Yoganarasimha et al. 2011, Hargreaves et al. 2005, Neunuebel et al. 2013). LEC has also been shown to be responsive during object-context association tasks (Wilson et al. 2013), and to display object-trace representations with cells’ firing fields forming at locations where an object had previously been placed (Deshmukh & Knierim 2013, Tsao et al. 2013). Evidence for temporal coding has been observed in both EC subregions, either throughout a trial at different task-relevant scales during context switching in LEC (Tsao et al. 2018) or during immobile waiting in MEC (Heys & Dombeck 2018, Heys et al. 2020). Despite the plethora of features encoded by functionally distinct cell types in EC, it’s not clear if all of these cell types participate in the direct projection to CA1 or to what degree the information about navigation, context, place or time could be reconciled from the activity observed at its termination.

There is accumulating evidence that EC circuits also participate in processing goal-related information (Sosa & Giocomo 2021). When presented with goal-related information, coherence between LEC and CA1 increases (Igarashi et al. 2014), implying that learned task parameters might be conveyed to the hippocampus from the EC. During cue-reward associations, dopaminergic input from the ventral tegmental nucleus has been shown to alter activity in superficial LEC based on reward expectation (Lee et al. 2021). Single cell recordings in human EC during a virtual reality (VR) task have also revealed a type of goal cells which were spatially selective relative to perceived goals and reactivate when participants had to remember the cued locations (Qasim et al. 2019). A dopaminergic modulation has not been reported in the MEC; however, recent studies of MEC revealed that both grid and non-grid spatial signals are modified by the presence of rewarded locations, suggesting a possible role for MEC in goal-directed navigation (Boccara et al. 2019, Butler et al. 2019). While the impact of goals on MEC tuning provides evidence counter to the notion that MEC representations are purely spatial, changing the context (visual cues) while animals run on a linear track elicited an almost complete remapping of the MEC spatial inputs to CA1, even though the goal location remained fixed (Cholvin et al. 2021). In CA1, context-independent reward tuning has been observed (Gauthier & Tank 2018) suggesting that other either intrinsic or extrinsic mechanisms besides just the MEC projection might be important to goal representations. Importantly, the relationship between context and goal representations has not been studied for the LEC projection; given the identified reward expectation tuning in LEC, it is possible that individual cells with consistent goal tuning between contexts would be observed in the LEC projection. Thus, the relative contribution of MEC and LEC inputs to providing CA1 goal-directed information remains unknown.

Consistent with the critical role of EC in navigation and associative learning, previous studies have demonstrated that lesioning or silencing EC disrupts internal representations of the external environment. Specifically, these studies identified deficits in path integration (MEC only) and associative learning, especially involving place-object relationships (both MEC and LEC) (Wilson et al. 2013, Roy et al. 2017, Kuruvilla & Ainge 2017). While recordings made from CA1 during these lesion studies provided some clues about the nature of the information conveyed along this pathway, they failed to provide a coherent picture. Chronic lesions of MEC are accompanied by an expansion of place field size, resulting in a reduction in the spatial information content of CA1 (Brun et al. 2008, Hales et al. 2014, Schlesiger et al. 2018), while transient, reversible manipulations of MEC networks can trigger remapping of hippocampal place fields (Miao et al. 2015, Kanter et al. 2017, Zutshi et al. 2022) but these effects are mixed (Rueckemann et al. 2016, Robinson et al. 2017). Older models produced place cell tuning by aggregating inputs from overlapping MEC grid cells of different phases and spatial scales (Samsonovich & McNaughton 1997, Bonnevie et al. 2013, Burgess et al. 2007, McNaughton et al. 2006, Fyhn et al. 2002). However, place cells persist in CA1 after pharmacological silencing of MEC grid cells (Brandon et al. 2014) or bilateral optogenetic silencing of MEC (Zutshi et al. 2022), demonstrating that place field formation is not necessarily dependant on grid cell activity. Even after a complete lesion of MEC, place fields persist and exhibit normal remapping (Schlesiger et al. 2018). In contrast to investigations of the MEC, few studies have directly addressed the influence of LEC on the spatial context of CA1 cells. There is also evidence that LEC lesions impair firing rate remapping (Lu et al. 2013); nonetheless, the full range of likely influence of the direct LEC inputs on CA1 as well as on CA1 place fields remains unclear. Studies performing inactivation or lesions of the EC have failed to find a clear dependency of CA1 tuning on the input from either EC subregion; instead these manipulations revealed only partial deficits in tuning indicating that perhaps this responsibility is shared between the EC subregions.

In summary, we still lack a understanding of circuit level mechanisms of direct cortical influences on hippocampal CA1 population dynamics, and how responsibilities are shared between EC subregions. Particularly, it would be imperative to characterize the specific tuning profiles maintained in the set of cells that project from the superficial EC to CA1. The vast majority of studies have focused on either the LEC or the MEC independently (Lee et al. 2021, Sasaki et al. 2015, Sosa & Giocomo 2021), rather then performing a comparison of the two regions under identical conditions. Additionally, especially when investigating the LEC, few studies have focused on goal-directed navigation, context change, and how spatial representations shift with navigational objectives (Save & Sargolini 2017). To bridge these knowledge gaps, we conducted a direct characterization of information conveyed by EC afferents to the hippocampus by performing two-photon Ca^2+^ imaging during a battery of VR navigation tasks. This experimental design gave us the ability to perform a direct comparison of the LEC and MEC inputs to CA1 during identical tasks as well as a high level of flexibility and control over the task parameters. Our study reveals both segregated and overlapping representations of task, location, and context in both EC projections, with navigational goals strongly biasing tuning of the LEC pathway, while MEC inputs demonstrate more specificity towards particular locations and contexts.

## Results

In order to selectively characterize the information conveyed directly from the EC to CA1, we performed *in vivo* 2-photon functional Ca^2+^ imaging of EC axons within *stratum lacunosum moleculare* (SLM) of CA1. In SLM, these direct EC inputs to CA1 are known to extensively form synapses with pyramidal cell distal dendrites as well as other local interneurons (Steward & Scoville 1976). During the imaging of axonal Ca^2+^ events, mice performed a series of virtual reality (VR) tasks designed to probe mnemonic and spatial information processing. Recombinant adeno-associated viral (rAAV-Synapsin-GCaMP6s) injections were targeted to either LEC or MEC, and expression confirmed via histological evaluation following the completion of experiments (Figure 1A-D). In order to perform the tasks, mice were head-fixed on a running wheel and traversed a linear environment in VR. The virtual environments used were 4m long and each traversal, or “lap” was followed by a 2s timeout, or inter-trial interval (ITI) when the VR screens remained dark. Following the ITI, mice were “teleported” back to the starting position (Figure 1E). We used `suite2p` to functionally detect regions of interest (ROIs), which corresponded to putative boutons or axonal segments, allowing us to record from 41,617 LEC ROIs from 32 FOVs and 34,604 MEC ROIs from 25 FOVs across all tasks (*n* = 10 mice LEC and *n* = 6 mice MEC, Figure S1). From these ROIs, we extracted fluorescence time series data and identified significant Ca^2+^ transients, which were used for subsequent analysis (Figure 1B, C, S1).

**Figure 1.**
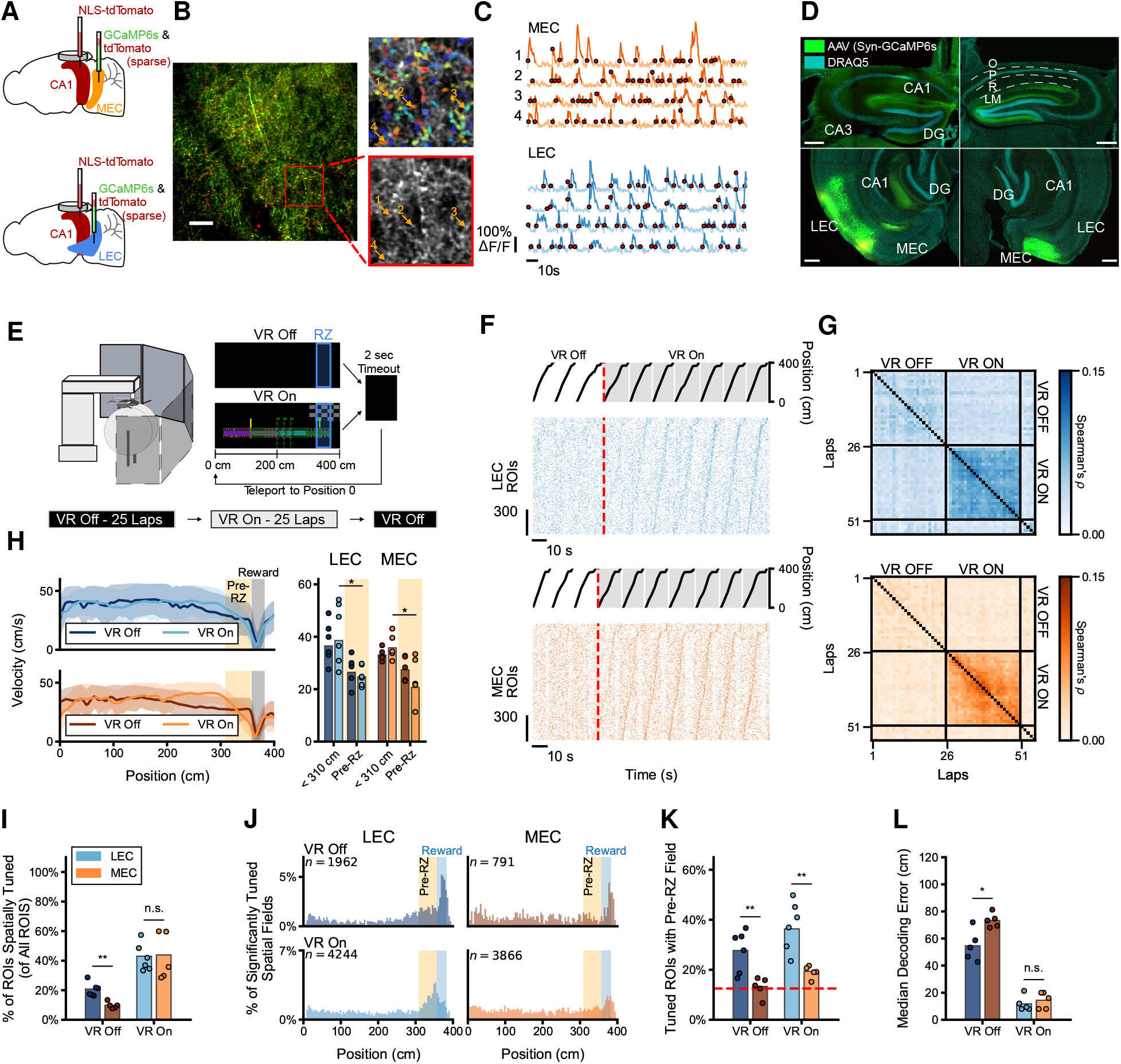
Visual VR is sufficient to invoke stable spatial tuning in both MEC and LEC axons. (**A**) Schematic of viral injections. Mice were injected with AAV1 Synapsin-GCaMP6s and sparsely expressing AAV9 Synapsin-tdTomato specifically in either the medial (top) or lateral (bottom) subregion of the entorhinal cortex. The sparse tdTomato in the EC axons was used as a fiducual for the motion correction algorthim as well as to assist in lining up the FOVs. An additional injection of diluted Nuclear Localized (NLS) Synapsin-tdTomato was targeted to the pyramidal layer of dorsal CA1 which was used to align the imaging FOV. (**B**) *Left*: Example time averaged FOV for imaging EC axons in SLM through intact hippocampus. Green is GCaMP6s, Red is tdTomato. Scale bar is 25*μ*m. *Right*: Magnified view of the indicated area identifying the axonal segments and boutons used to extract signals. Both the time average from the recording (*bottom*) and the extracted spatial masks (top) are shown. Example shown are MEC axons, for LEC example FOV see Figure S1D. (**C**) Example Δ*F/F* and detected transient Ca^2+^ events for both LEC (top) and MEC (*bottom*) axons. The numbered traces correspond to the numbered ROIs in B. Transient events indicated in bold and start times shown by the red circles. (**D**) Specificity of injections was confirmed for LEC (*left*) and MEC (*right*) injections following experiments via confocal imaging of sections. Both coronal sections of the hippocampus (top) from beneath the imaging window and a horizontal sections of EC (*bottom*) to confirm injection location are shown. Scale bars indicate 350 *μ*m. (**E**) Mice were trained to run while head fixed on a running wheel while VR cues were displayed on 5 LCD monitors surrounding the animals. Water deprived mice ran for 25 laps with no visual cues and one fixed reward location every 4m lap, followed by 25 laps with visual cues and the same reward location. Reward delivery was a non-operant water drop presented to animals when they reached the reeward zone, however mice which stopped and continued to lick would receive additional water for up to 2 seconds. After each lap there was a 2s timeout when the VR was dark after which the animals were teleported back to position 0. (**F**) Position of an animal along the 4m lap and accompanying raster plot of Ca^2+^ event start times during the last 3 VR Off laps and first 7 VR On laps for both LEC (top) and MEC (*bottom*). ROIs sorted by peak firing location during the VR On laps. Note that sequences emerge rapidly after exposure to spatially fixed visual cues for both the LEC and MEC labeled axons. (**G**) Spearman’s rank correlation coefficient (*ρ*) calculated between each pair of laps on the population vector of tuning curves for all ROIs at every position averaged across mice for both LEC labeled FOVs (top) and MEC (*bottom*). (**H**) Velocity during goal directed navigational task. *Left*. Mean velocity at each position for both the VR Off and VR On conditions, shaded region depicts standard deviation between mice. *Right*. Quantification, by mouse of the mean velocity far from the RZ (position < 310 cm) and close to the RZ (position 310 - 360 cm). There was a significant impact of RZ distance (near vs. far), but not EC subregion or VR condition. Linear mixed-effects model with mouse as the random factor and EC subregion, VR condition and RZ distance as fixed factors, *n* =6 mice LEC, *n* = 5 mice MEC, effect of RZ distance: *p* < 0.001, EC subregion: *p* = 0.228, VR condition: *p* = 0.346. For both LEC and MEC labeled mice, mean velocity significantly dropped just prior to RZ. Two-sided paired t-test, LEC: *t* = 3.1622, *p* = 0.0242, MEC: *t* = 3.1619, *p* = 0.0341, and, during the VR Off condition, there was a noticeable, but not significant trend for the mice to slow down in the 50cm prior to the RZ. Two-sided paired t-test, LEC: *t* = 2.2538, *p* = 0.0739, MEC: *t* = 2.6917, *p* = 0.0546. (**I**) More axons were determined to be significantly spatially tuned (*p* < 0.05) during VR On as opposed to VR Off conditions. LEC: VR Off 20.8%, VR On 43.0%, *n* = 6 mice, two-sided paired t-test *t* = 5.6881, *p* = 0.0023. MEC: VR Off 9.8%, VR On 43.8%, *n* = 5 mice, *t* = 4.9411, *p* = 0.0078. Additionally, LEC had significantly more tuned ROIs then MEC without visual cues, but not during the VR On laps. Welch’s unequal variances t-test VR Off: *t* = 4.9666, *p* = 0.0013, VR On: *t* = 0.0720, *p* = 0.9448. Significant effect of VR condition (On vs Off), two-factor ANOVA, *F* = 45.7002, *p* = 2.468*e* - 6, but not axon type *F* = 1.6117, *p* = 0.2204, or the interaction between type and condition, *F* = 2.0007, *p* = 1.7429, (**J**) Percentage of all spatially tuned fields with a centroid at each location. An enrichment of LEC axons with spatial fields in the 50 cm area prior to the start of the reward zone (highlighted). *Top*. VR Off condition LEC: *n* = 1962 spatial fields, MEC: *n* = 791 spatial fields. *Bottom*. VR On LEC: *n* = 4244 spatial fields, MEC: *n* = 3866 spatial fields. (**K**) A higher percentage of stable spatial fields were found in the 50 cm prior to the reward zone in the LEC axons as opposed to the MEC in both conditions, *n* = 6 mice LEC, *n* = 5 mice MEC, VR Off - LEC 27%, MEC 13%, *p* = 0.0060, VR On - LEC 36%, MEC 19%, *p* = 0.0049, Welch’s unequal variances t-test. Significant effect of both axon type and VR condition (On vs Off), two-factor ANOVA, effect of axon type *F* = 29.2826, *p* = 3.845*e* – 5, VR condition *F* = 7.2443, *p* = 0.01492, interaction of type and VR condition (not significant) *F* = 0.4167, *p* = 0.5267. (**L**) Decoding location from the activity of LEC and MEC axon ROIs was equally effective. Median error decoding of animal’s virtual location was calculated for 1000 randomly selected ROIs per mouse, 3 different randomly selected splits where used and results averaged. LEC: mean error VR Off 54.4 cm, VR On 11.7 cm, *n* = 5 mice, two sided paired t-test, *t* = 11.6847, *p* = 0.0003, MEC: VR Off 73.3 cm, VR On 14.4 cm, *t* = 16.4268, *p* = 0.001. Position decoding was also significantly better for LEC then MEC ROIs during the VR Off condition but not for the VR On. Welch’s unequal variances t-test, VR Off: *t* = 3.2979, *p* = 0.0185, VR On: *t* = 0.3367, *p* = 0.4912. Significant effect of axon type, VR condition, and the interaction between type and condition, two-factor ANOVA effect of axon type *F* = 10.0078, *p* = 0.0060, VR condition *F* = 228.1845, *p* = 6.876*e* – 11, interaction of type and VR condition *F* = 5.6294, *p* = 0.0353.

### Stable representations of space in both LEC and MEC during 1D goal directed navigation task

Given the rich spatial and sensory responses previously identified in local recordings of the superficial EC (Deshmukh & Knierim 2011, Sasaki et al. 2015), we first investigated how a virtual environment was encoded in the activity of the direct EC-CA1 projection using our axonal imaging method. We used a spatial goal learning task where mice first ran for 25 laps without access to any visual cues (i.e. in the dark, “VR Off” condition), followed by 25 laps with cues (“VR On”, Figure 1E). Both conditions featured a water reward located near the end of the 4m lap. As expected, the animals were able to anticipate the presence of the reward better when they had access to the visual cues, indicated both by a reduced running velocity and increased rate of anticipatory licking when they approached the rewarded location. Mice also exhibited these behaviors when there was no access to the VR cues, albeit with substantially less accuracy, presumably through path integration between visits to the rewarded location (Figure 1H, S3A).

In order to determine if there was any underlying subregion or cue related differences in the signals we recorded, we examined the properties of the Ca^2+^ events in each context. In both VR Off and VR On conditions, for both LEC- and MEC animals we observed transient Ca^2+^ events of similar duration, height, and rate (Figure S2A-C). Since mice clearly took advantage of the VR cues to more accurately navigate, we next asked how the presence or absence of the VR environment impacted the activity of EC axons. To answer this, we investigated the similarity between laps of the Ca^2+^ events (Figure 1F), constructing a cross correlation matrix for the population vector (PV) of all the tuning curves for all axons in each field of view (FOV). Access to the visual cues led to more stable representations of the virtual space existed in both projections to CA1. Correlations for spatial PVs were significantly higher between all pairs of VR On laps, compared to VR Off, for both LEC- and MEC-expressing cohorts (*p* = 1.869 × 10^-7^ effect of VR condition, two factor ANOVA, Figure 1G, S3B). Additionally, during the VR On condition there was no significant difference between the lap-to-lap correlations between the LEC and MEC axons (*p* = 0.3021 unequal variances t-test, Figure S3B). The increase in the correlations appeared within the first laps following the introduction of the spatial cues and persisted for the duration of the subsequent 25 laps in which animals had access to the cues, indicating that stable contextual representations in the EC may form quickly and then maintain their tuning across many subsequent exposures (Figure 1G). This finding that EC axons quickly form stable representations of virtual environments is consistent with previous reports of MEC axons (Cholvin et al. 2021) and expands this to the LEC projection.

We additionally evaluated the tuning of individual ROIs and similarly found increased spatial tuning in both the LEC and MEC axons during the VR On condition compared to the VR Off (Figure 1I, S3C). Compared to shuffled data 43% of the LEC ROIs and 44% of the MEC ROIs were significantly spatially tuned (*p* < 0.05) when the animal had access to visual cues. Interestingly, without the visual cues, 21% of LEC ROIs still were spatially tuned while only 9% of the recorded MEC ROIs were. This both confirms that the visual cues elicited significantly more spatially stable responses in both LEC and MEC axonal projections, but also highlights a task-dependant dissociation between the LEC and MEC activity (Figure 1I). For the spatially tuned axons, there was a significant trend towards increased reliability and specificity during the laps in which the visual cues were present, as well as increased field width in the MEC axons (Figure S3D-F). In order to ensure that there were no systematic influences of imaging artifacts that could explain the observed differences between the MEC and LEC cohorts, we calculated the imaging frame stability (after motion correction) and found no trend between average frame correlation and percentage of spatial fields–nor was there any significant difference in the frame stability metric between the LEC and MEC FOVs (Figure S3G).

It was unexpected that such a large proportion of the LEC axons would exhibit significant spatial tuning, even in the absence of visual cues (Figure 1I, S3C). In particular, we also saw a notable enrichment of the LEC spatial fields just prior (50 cm) to the reward zone in both conditions, that was much greater than any clustering of spatial fields in the MEC axons (percent of spatial fields within 50 cm of the reward location, VR Off LEC: 27.7%, MEC: 13.3%, VR On LEC: 36.4%, MEC: 19.4%, Figure 1K). Interestingly, the set of axons participating in the enrichment around the RZ showed significantly more overlap for the LEC axons, as compared to the MEC (Figure S3I). When the VR cues were present, reward proximity/anticipation tuning of the LEC axons was observed in all LEC FOVs (Figure 1K) and the degree of the enrichment was significantly correlated with the animal’s anticipatory licking behavior (Figure S3H). In the VR Off condition there was also increased spatial tuning within and immediately following the reward zone, which was less surprising given that the reward delivery was the only spatially fixed cue during these laps and a similar confluence of spatial fields was also observed in the MEC axons (Figure 1J). These results demonstrate a goal anticipatory bias in the tuning of the LEC activity, which could be partially distinct from the presence (or absence) of the current contextual cues, while the spatial activity in the MEC axons might be, to a much higher degree, dependant on the VR environment.

The observation that LEC axons projecting to CA1 were spatially tuned, in some cases just as much or more than axons originating in the MEC, is in stark contrast to studies which have sought to identify the foundations of the hippocampal place code purely targeting MEC cell populations (Zhang et al. 2014, Cholvin et al. 2021). We therefore sought to quantify the degree to which the input from each EC subregion was independently mapping the virtual environment. Using a linear SVM to decode the location of the mice (Stefanini et al. 2020), decoders trained on both LEC and MEC axonal activity performed equally well during the “VR On” condition (cross-validated median decoding error, mean by mouse LEC: 11.7 cm, MEC: 14.4 cm, *p* = 0.4912 unequal variances t-test, Figure 1L). Additionally, both LEC and MEC classifiers had significantly better performance when the animals had access to VR cues than when they did not (VR Off decoding median error, LEC: 54.4 cm, MEC: 73.3 cm, Figure 1L-D). Increasing the number of ROIs used for decoding improved the decoding accuracy, but this improvement plateaued after 500 - 750 ROIs (Figure S4A). For this reason we used identical numbers of ROIs per FOV/mouse when performing any decoding analyses. Despite the enrichment of tuned spatial fields specifically found in the LEC projection near the reward, position could be accurately predicted from both the LEC and MEC axons throughout the entire VR track (Figure S4B, C), suggesting that during goal directed navigation LEC also contributes to construction and refinement of hippocampal spatial maps and that this contribution is not necessarily restricted to areas near the reward, but distributed throughout the entire 4m environment. When visual cues were not provided, the activity in the LEC axons was actually more informative about the animal’s location when the animal is near to the reward zone, whereas using the MEC axons, the decoding error remains, on average quite high (LEC median error, VR Off, reward zone far 67.3 cm, reward zone near 35.1 cm, MEC reward zone far 83.9 cm, reward zone near 58.2 cm, Figure S4C). Notably, when the VR cues are off, the reward was the only spatial cue that animals had access to and the decoding based on the LEC axons outperforms decoding on the MEC axons (Figure 1L, S4D). This observation that decoding position from the LEC axons during the VR Off condition is in agreement with the observed larger proportion of spatially tuned axons and, as expected, we find better decoding performance is related to the fraction of spatially tuned axons (Figure S4E, F).

Given previous reports of velocity modulation of MEC neurons (Hardcastle et al. 2017, Kropff et al. 2015) we finally asked if it was possible to decode the animals’ velocity based on the activity of the cell population that projection to CA1. Across mice, we found that it was, in fact, possible to decode velocity from the LEC axonal activity, and there was a non-significant trend for successful decoding in the recorded MEC axons during the VR On laps (Figure S4G). Thus, it is unlikely that a large proportion of MEC “speed cells” (or at least unlikely that a large proportion of cells purely selective for velocity) project to CA1–a conclusion in agreement with previous observations of velocity modulated cells being predominantly in MEC layer II, with only a lesser (although not insignificant) degree of velocity modulation of cells in layer III (Kropff et al. 2015).

### LEC-CA1 projection transmits a reward-centric map of space during a goal alternation task

The prominent enrichment in LEC projection in anticipation of fixed reward locations suggests that a large portion of the spatial stability in the activity of the LEC projection might be due to the presence of a consistent, familiar, goal location, while the MEC projection might more uniformly represent locations in the virtual context. In order to test this directly, we next compared the activity of each EC projection while the animals were forced to adapt to changes in the rewarded location. To this end, we recorded from EC axons while mice performed a reward location alternation task, in which the location of the water reward would alternate every 10 laps between one of two fixed locations (RZ1 and RZ2, Figure 2A). The visual cues remained constant between reward-location blocks and there were no cues that the goal change occurred, other than the presentation of the reward itself. Mice were familiar with the task prior to the imaging session, but had to learn to adjust their behavior following the change in reward location during the imaging session. Both LEC and MEC cohorts of mice learned the reward locations and followed the change within a lap of the switch, shown by the anticipatory licking which shifted markedly toward the new location following reward translocation (Figure 2B). Mice also tended to run at a similar velocity along the entire track at all locations, except for those adjacent to the reward zone and we observed no significant differences in the animals’ average running velocity between reward locations (Figure S5A).

**Figure 2.**
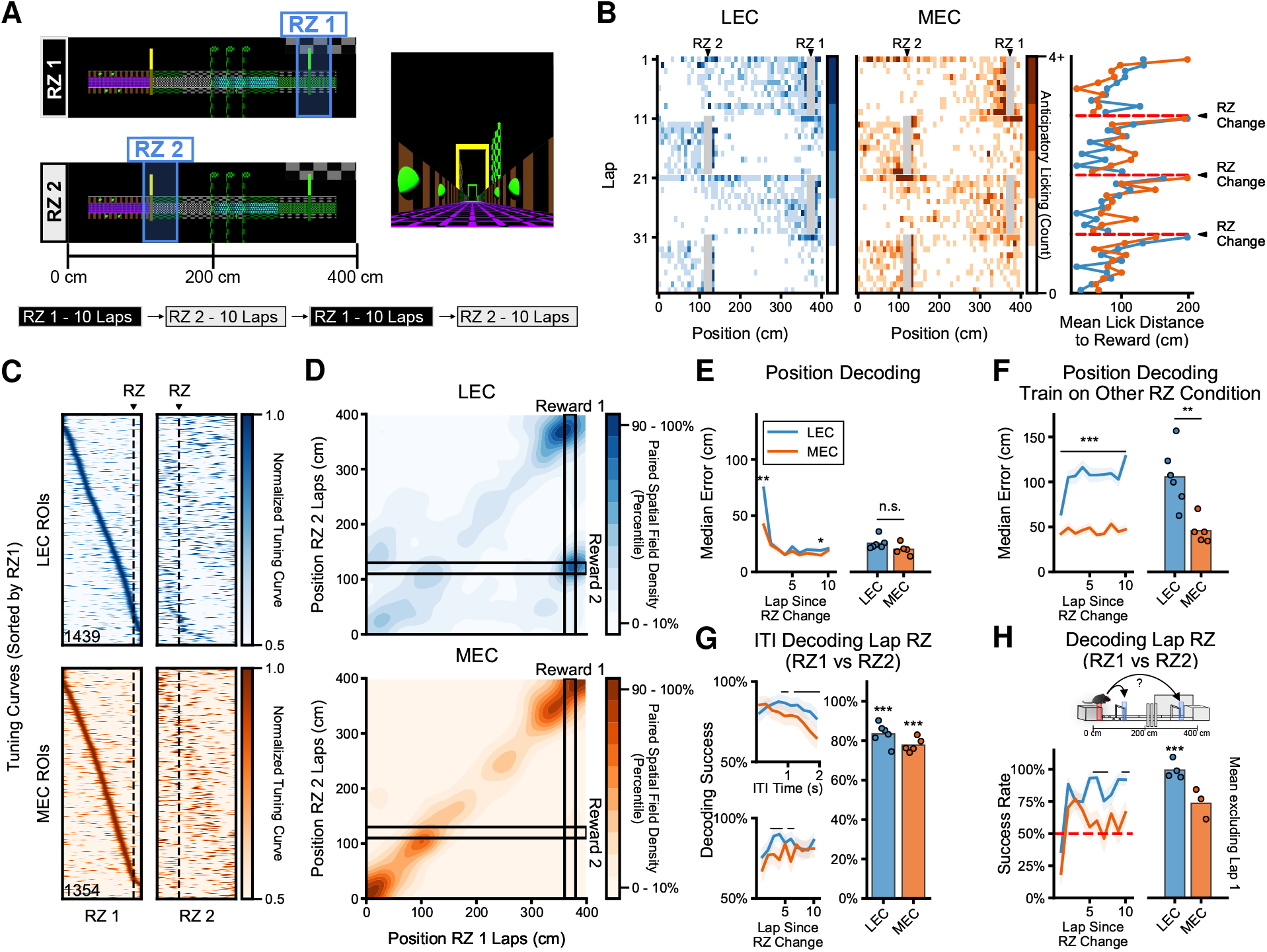
LEC axons activity reflects a change in goal location while MEC axons remain spatially stable. (**A**) Diagram of the reward alternation task. Mice learned two different reward locations, with the reward location moving between them every 10 laps. (**B**) Heat maps showing the locations where mice licked at the reward valve in anticipation of the reward zone. Both cohorts of mice with either LEC (*left*) or MEC (*center*) axons labeled learned the new goal locations throughout the 10 lap period as evident by anticipatory licking. *Right*. Mean distance from the anticipatory licks to the RZ always increases on the laps immediately following the change. (**C**) Tuning curves for all ROIs in a single FOV of LEC axons (Top) and MEC axons (*bottom*) recorded during the alternating reward task. ROIs sorted by the peak Ca^2+^ event rate across all RZ1 laps. (**D**) Between RZ condition paired spatial field density, centroid locations calculated during RZ1 laps (x-axis) compared to RZ2 laps (y-axis). Spatially tuned LEC (top) ROIs showed more remapping then MEC (*bottom*) following RZ change and several follow reward location. (**E**) Position was decoded from the activity of 900 randomly selected ROIs per mouse from both the LEC and MEC axons. *Left*. Similar median error per lap except for the first lap when decoding from LEC axons was significantly worse. Welch’s unequal variances t-test, 3 randomly selected sets of ROIs per mouse and 2 × 2 RZ exposures, LEC: *n* = 36, MEC: *n* = 30, Lap 1: *p* = 0.0027. *Right*. Average across all laps there was no difference between the LEC and MEC mice. Welch’s unequal variances t-test, LEC: *n* = 6 mice, MEC: *n* = 5 mice, *p* = 0.4421. (**F**) Decoding the position on the alternate RZ laps resulted in significantly worse median error per lap for the LEC axons compared to the MEC. *Left*. LEC worse then MEC at all laps within the block of trials. Welch’s unequal variances t-test 3 randomly selected sets of ROIs per mouse and 2 RZ exposures LEC: *n* = 36, MEC: *n* = 30, *p* ≤ 0.003 for all laps. *Right*. The average by mouse across all laps was also significantly worse. Welch’s unequal variances t-test, LEC: *n* = 6 mice, MEC: *n* = 5 mice, *t* = 4.0507, *p* = 0.0047. (**G**) The current location of the reward was decoded during the 2 second inter-trial interval. *Left-Top*. Classification of the RZ was successful throughout the inter-trial interval and (*Left-Bottom*) at all laps within the RZ block. Significant differences between the LEC and MEC axon ROIs indicated by black line. Welch’s unequal variances t-test, LEC: *n* = 6 mice, MEC: *n* = 5 mice, *p* ≤ 0.05. *Right*. Averaged by mouse, decoding RZ during the ITI better then 50% for both LEC and MEC data. One-sided one-sample t-test, LEC: *t* = 7.7194, *p* = 1.0*e* – 5, MEC: *t* = 8.7109, *p* = 3.2*e* – 5. Decoding performance on average was not significantly better for LEC or MEC, by mouse average performance LEC: 83.5%, MEC 77.8%, not significant, Welch’s unequal variances t-test *t* = 2.1163, *p* = 0.0645. (**H**) The current location of the reward was decoded at a fixed location during the initial portion of the track. Velocity was binned (10 cm/s) and matched between RZ1 and RZ2 laps, and frames when the animal was licking were excluded. *Left*. Classification of the RZ was successful on more laps for the LEC axons then the MEC axons. Significant differences between the LEC and MEC axon ROIs indicated by black line, *p* ≤ 0.05, LEC: *n* = 4 mice, MEC: *n* = 3 mice, Welch’s unequal variances t-test. *Right*. Averaged by mouse, decoding RZ better then 50% for the LEC ROIs but not MEC. One-sided one-sample t-test, LEC: *t* = 6.0396, *p* = 0.0006, MEC: *t* = 1.1802, *p* = 0.0711

Despite exhibiting similar performance on the behavioral indicators of task competency (i.e. running and anticipatory licking), differences in the tuning of the EC axons between reward conditions were drastic. LEC tuning considerably reorganized upon RZ translocation, while the MEC projection remained more stable (Figure 2C). The greater continuity of the spatial representations in the MEC data between RZ1 and RZ2 blocks indicates that MEC activity was more influenced by the VR/spatial cues than location of the reward (Figure S5B). For both the LEC and MEC axonal FOVs, we observe a similar percentage of spatially tuned ROIs during the RZ1 and RZ2 laps (34 - 42%, Figure S5C); additionally, about 30% of both the LEC and the MEC ROIs that were stable in one reward zone condition were stable in both (Figure S5D). When we looked at the spatial stability of these ROIs, however, the locations of the spatial fields of the MEC axons shifted less between reward zone conditions compared to those of the LEC axons (Figure S5D). Strikingly, there was a population of LEC ROIs that followed changes in the reward location vicinity, resulting in a greater overlap of in axons with spatially tuned fields in areas near the rewarded location compared to the MEC (mean Jaccard overlap score for reward tuned ROIs, LEC: 0.2214, MEC: 0.0721, *p* = 0.0137 unequal variances t-test, Figure 2D, S5E).

We next asked how the reward change triggered remapping in axonal tuning impacted our ability to predict the animals’ location from the activity of the axons. Interestingly, LEC activity was significantly better at predicting the animal’s position on laps when the reward location was consistent and likely known by the animal. The LEC had notably worse performance at decoding space on the first lap following the reward zone switch, but otherwise has similar accuracy to the MEC (mean by mouse first lap error LEC: 74.67 cm, MEC: error 41.8 cm second lap error LEC: 25.4 cm, MEC 23.9 cm Figure 2E). Additionally, training the decoder on laps from the opposite reward condition yielded worse decoding performance from the LEC than from the MEC (average of median decoding error by mouse LEC: 106.5 cm, MEC: 45.8 cm, Figure 2F), suggesting that the LEC had significantly shifted representations of the virtual environment. This reward translocation-related reorganization only took 1 lap following the reward location had change, after which the decoder performance rapidly plateaued, indicating that spatial representations in the LEC projections reorganize quickly and have separable representations of the current reward zone condition.

Since the tuning of the LEC axons switched between distinct representations of the same virtual track upon subsequent reward translocations, we asked if there was an internal representation of the current task demands–asking is the current reward zone location between laps preserved within the EC between laps? We therefore looked to see if there was a representation of the current reward zone location in the EC during the 2s inter-task interval, when the VR is off. We found it possible to decode the current reward zone location from EC axonal activity throughout the timeout, and that this was successful for all laps except the first lap of a reward zone context block (Figure 2G). The previous lap reward location could be decoded from both then LEC and MEC projections to CA1 better than chance and, although the average decoder performance from the LEC axons was higher (by mouse average performance LEC: 83.5%, MEC 77.8%) this difference was not significant (Figure 2G). Importantly, there was no reward location dependent difference in the animals’ running or licking behavior during the timeout: the animals did not lick and ran at similar velocities regardless of the current reward zone (Figure, S5F). Similarly, we found that, for matched location, velocity and (no) licking epochs, it was possible to decode the current reward zone location from the activity of the LEC axons significantly better than chance after the first lap in the reward zone block (Figure 2H). Together, these results indicate that the EC inputs to CA1 contain information relevant to navigational variables that extend beyond the current position of the animals or cues on the VR screens. Rather decoding a reliable “spatial” map from the LEC is possible, but dependent on an understanding of the current navigational task. This implies a degree of prospective or retrospective coding (Frank et al. 2000) may be occurring, which could have ramifications for the animal’s route planning, since, importantly, this information could be resolved to a “reward-vector” (similar to landmark vector tuning previously described in Deshmukh & Knierim 2013), tracking the distance to the next reward or from the previous one.

In order to further dissect the relationship between the reward zone location and the dissociation between LEC and MEC spatial representations, we performed a modification of the alternating reward task. For the modified task, we introduced a block of probabilistic reward delivery. During which on 50% of the laps no reward was issued (Figure 3A). In this way, we could compare the peri-reward zone spatial tuning enrichment in EC axons with the animal’s expectation of reward. Using trial blocks when reward was reliably present as a comparison, we found that the MEC representations remained spatially stable throughout the experiment (regardless of reward location or whether the reward was guaranteed or stochastic). However, the LEC tuning curves were robustly altered by the changes to the rules governing reward delivery (Figure 3B, C). Specifically, during the block of probabilistic reward laps, we saw consistent tuning before the expected reward location; then, on the subset of laps when the reward was omitted, representations diverged and the remainder of the lap was no longer similar to the reliable RZ2 laps (Figure 3D). This difference was observed comparing the PV correlation of the tuning curves before and after the RZ2 location (positions 110-130 cm, Figure 3E). Comparing changes in the PV of the LEC axonal activity around the two reward zones indicates that the animals’ expectation of reward delivery significantly affects the activity of the LEC axons, but no relationship like this was found for the MEC labeled axons. These results demonstrate the reliance on an expected goal location for the LEC axons to generate spatially informative maps. The LEC axons maintain the spatially stable representation until the reward is unexpectedly omitted, the resulting reorganization of the activity in the LEC axons is notably distinct from what is observed in the MEC axons, which maintain more stable spatial tuning curves PVs regardless of reward delivery or omission.

**Figure 3.**
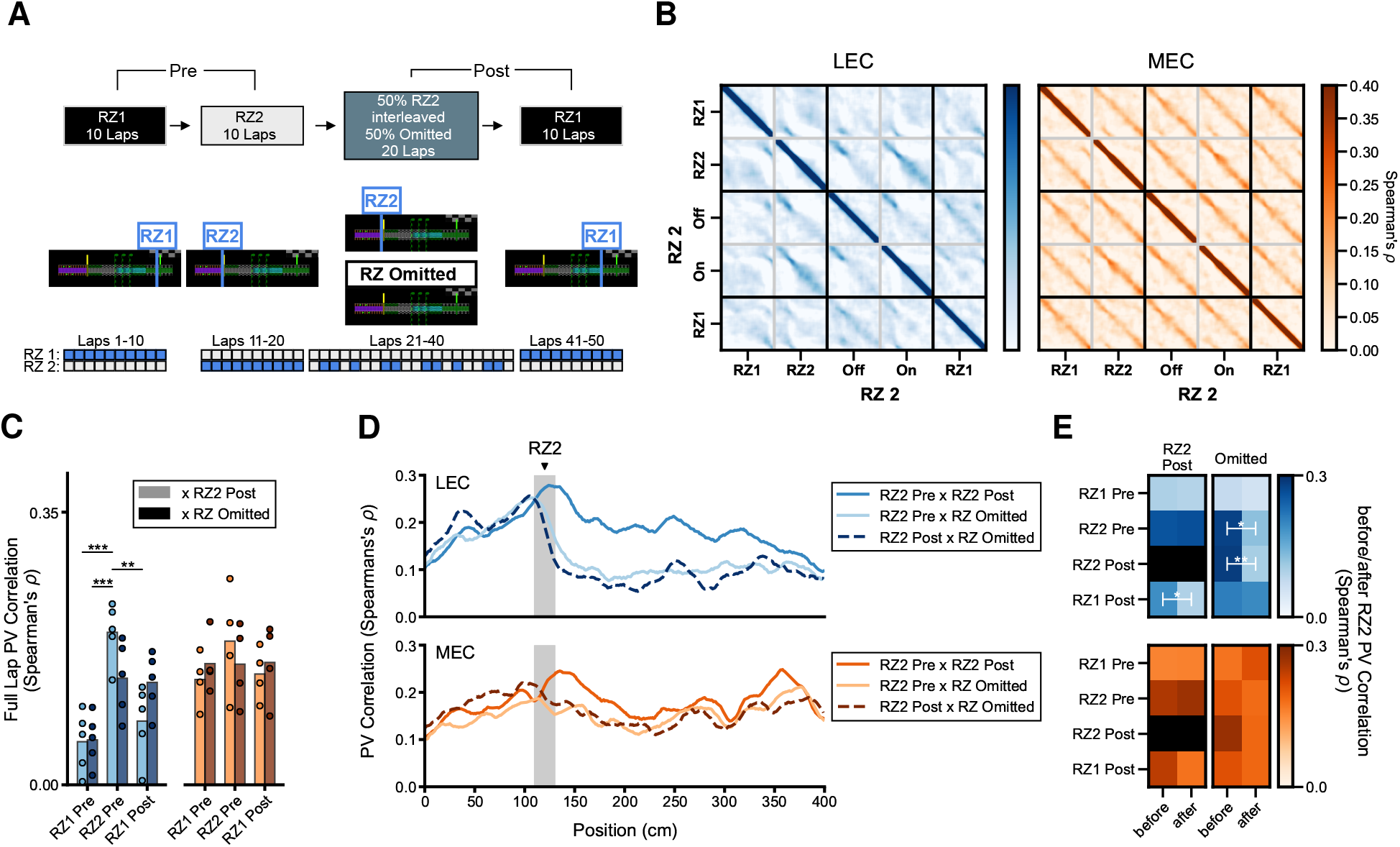
LEC axons activity reflects a change in goal location while MEC axons remain spatially stable. (**A**) Schematic of the modified task structure with probabilistic rewards block. *Bottom*. Reward delivery schedule, blue boxes indicate rewarde laps, top row RZ1, bottom row RZ2. (**B**) Correlation (Spearman’s *ρ*) between population vectors of mean Ca^2+^ event rate for every position for each RZ block of the experiment (*Left*. LEC, *Right*. MEC). RZ2 On and Off trials were interleaved in the third block of the experiment are listed separately here (see **A**). MEC tuning was more spatially stable regardless of current reward schedule, while peri-RZ representations shift in LEC representations, Mean across LEC: *n* = 5 mice, MEC: *n* = 4 mice. (**C**). PV correlations during probabilistic reward delivery with all other blocks. When the reward was delivered, the LEC activity was significantly more correlated to the RZ2 template, then the other trial blocks. When the reward was omitted, PV correlation to RZ2 and RZ1 (Post) blocks were similar. Two-factor repeated measures ANOVA, LEC: *n* = 5 mice, effect of template *p* < 0.0001, interaction of template and reward presence *p* = 0.0166, MEC: *n* = 4 mice, effect of template *p* = 0.0785, interaction of template and reward presence *p* = 0.4623. (**D**). The LEC representation of the track is more dependant on the presence or absence of reward than MEC. PV correlation by position between RZ2 (Pre or Post) and RZ Omitted laps drops after the expected RZ. (**E**) The PV correlation in the 50 cm prior to RZ2 compared with the 50 cm after (highlighted locations in E). The LEC representations become dissimilar dependant on whither the reward was delivered (RZ2 Pre/Post) or not (RZ Omitted), but not MEC. PV correlation (Spearman’s *ρ*). * - *p* < 0.05, ** - *p* < 0.01.

### MEC axons remap following context change, while LEC axonal representations are more stable with fixed goal location

In the previous section, we showed that rewarded locations and reward expectation strongly influence LEC-CA1 axonal activity. However, we have not yet considered the role of context in EC axonal representations. Since CA1 has been shown to remap in response to changes in context (Colgin et al. 2008, Leutgeb et al. 2004, 2005, Frank et al. 2004), we next asked how the EC-CA1 projection reorganize their activity if we completely alter the visual scenery while leaving the reward zone at a fixed location (Priestley et al. 2022). Similar to the block trial structure used in the reward translocation task, we exposed the animal to two different VR contexts in alternating blocks of 10 trials (Figure 4A, S5A). Sorting MEC axonal Ca^2+^ traces by their activity during Context A laps revealed that not much structure remained visible during Context B, interestingly some spatial tuning remained apparent in the LEC axons, likely influenced by the fixed reward location (Figure 4B). We did observe that during laps through the same context, the ROIs had more similar spatial tuning curves than when comparing across contexts for both MEC and LEC subregions (Figure 4C, D, S6C). Although, LEC axonal activity is more correlated between different contexts than the MEC activity–in particular, LEC axon responses are most strongly correlated at locations directly adjacent to the reward location (Figure 4D). Thus the fixed reward location relative to the start/end of the lap results in a more similar spatial stability in LEC despite the change of the VR context.

**Figure 4.**
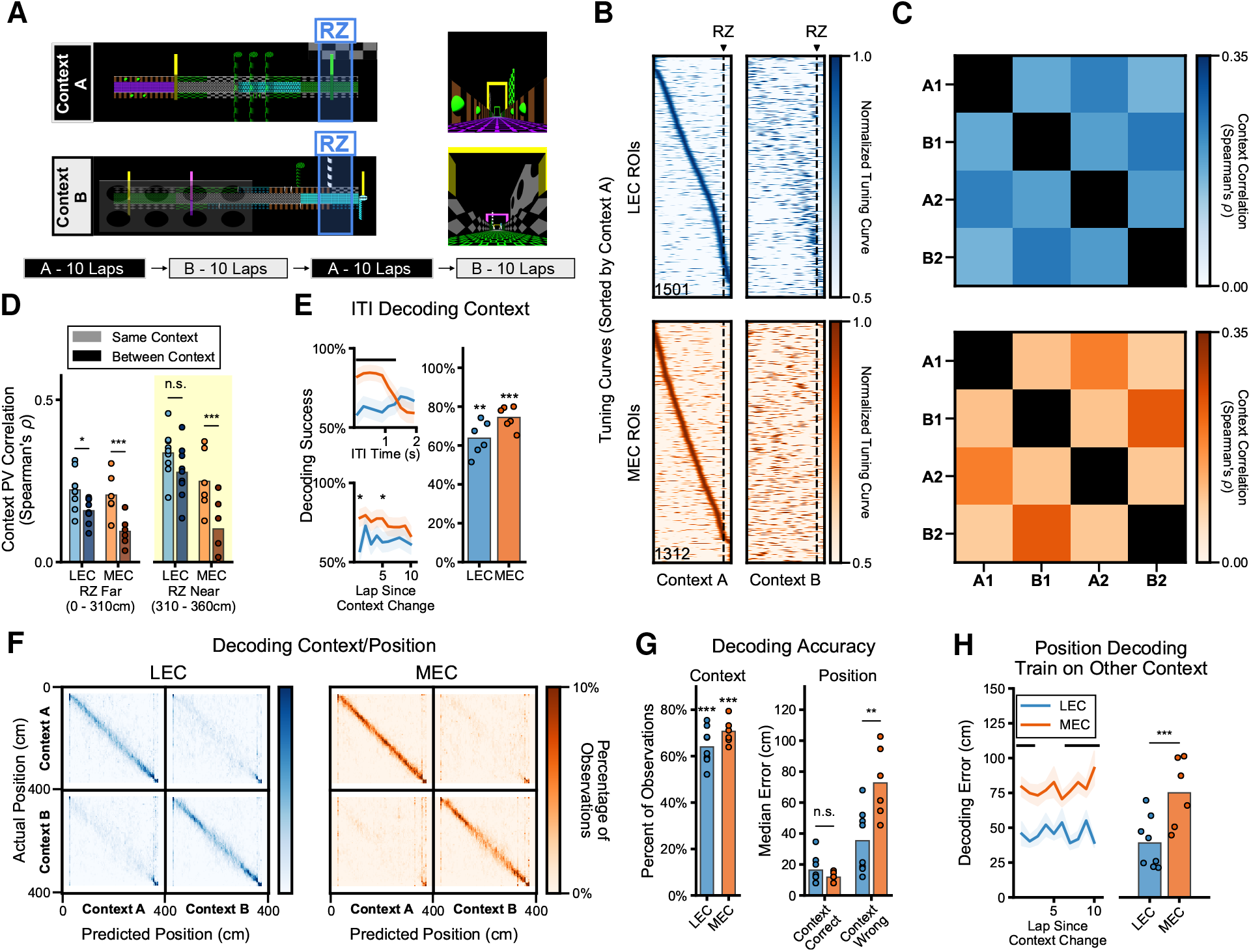
MEC responses remap more significantly during context change while LEC shows consistent tuning with respect to track start and reward. (**A**) Diagram of the context alternation task. The mouse was presented with two distinct visual virtual reality contexts, A and B which alternate every 10 trials, for 40 trials. (**B**) Tuning curves for all ROIs in a single FOV of LEC axons (Top) and MEC axons (*bottom*) recorded during the alternating context task. ROIs sorted by the peak Ca^2+^ event rate across all Context A laps. (**C**) Spearman’s rank correlation coefficient between population vectors calculated individually per block of 10 laps in each context. Mean across mice, LEC: *n* = 9 mice, MEC: *n* =6 mice. (**D**) Comparison of same context (A-A or B-B) and different context (A-B) PV correlation (Spearman’s *ρ*). PV correlations were higher near RZ (310 - 360 cm) in the LEC axons as compared to the rest of the lap (< 310 cm), or the MEC. Linear mixed effects-model with RZ distance (near vs far), comparison type (same vs different) and axon type (LEC vs MEC) as fixed factors and mouse as the random factor, LEC: *n* =9 mice, MEC: *n* =6 mice, 2 same context comparisons and 4 different context comparisons per mouse, and 2 RZ distances: axon type factor *z* = 7.249, *p* = 0.0642, context comparison type *z* = 5.387, *p* = 7.160*e* – 8, RZ distance 12.174, *p* = 4.288*e* – 34, axon type x RZ distance interaction *z* = 7.209, *p* = 5.619*e* – 13, axon type x comparison type interaction *z* = 2.527, *p* = 0.0150, RZ distance x comparison type interaction *z* = 0.268, *p* = 0.7890, RZ distance x comparison type x axon type *z* = 1.478, *p* = 0.1394. Tukey test for multiple comparison of means: * - *p* < 0.05, ** - *p* < 0.01, *** - *p* ≤ 0.001. (**E**) The current context decoded during the 2 second ITI. *Left-Top*. Classification of the context was successful throughout the inter-trial interval. *Left-Bottom*. Classification was also successful at all laps within the context block. Significant differences between the LEC axons and the MEC indicated by black line. Welch’s unequal variances t-test, LEC *n* = 6 mice, MEC: *n* = 5 mice, *p* ≤ 0.05. *Right*. Averaged by mouse, decoding context during the ITI better for MEC than LEC ROIs. Welch’s unequal variances t-test, LEC: *n* = 6 mice, MEC: *n* = 6 mice, *t* = 2.4993, *p* = 0.0347. Also better than chance (50%) for both LEC and MEC ROIs. One sided one sample t-test LEC: *t* = 1.9346, *p* = 0.0059, MEC: *t* = 5.1634, *p* = 0.0001. (**F**) Confusion matrices for decoding position and context from the EC axon ROI activity from 750 randomly selected ROIs per mouse. (**G**) *Left*. Both LEC and MEC ROIs predicted the correct context significantly higher then chance levels. One sided, one sample t-test on percent correct by mouse, LEC: *n* = 8 mice, *t* = 5.1472, *p* = 0.0007, MEC: *n* = 6 mice, *t* = 9.1988, *p* = 0.0001. *Right*. Position decoding errors were similar between LEC and MEC ROIs when the context was correctly decoded, however decoding position from the LEC mice was much more accurate when the context was incorrectly identified. Welch’s unequal variance t-test, LEC: *n* = 8 mice, MEC: *n* = 6 mice, *t* = 3.1611, *p* = 0.0099 for incorrect context, *t* = 1.3098, *p* = 0.2775 when context was decoded correctly. (**H**) *Left*. Decoding the position from the axon activity from 750 ROIs recorded on the alternate context laps results in significantly worse median error per lap for the MEC axons compared to the LEC axons at most within the block of trials. Welch’s unequal variances t-test, 3 randomly selected sets of ROIs per mouse and 2 context exposures LEC: *n* = 36, MEC *n* = 30, *p* ≤ 0.003 for all laps. *Right*. The average by mouse across all laps was also significantly worse for MEC axons compared to LEC. Welch’s unequal variances t-test, LEC: *n* = 6, MEC: *n* = 5, *p* = 0.0047.

As with the other tasks, we calculated spatial tuning scores for every ROI imaged during this task in order to look at spatial selectivity and remapping on at the individual axon level. While the LEC axons showed similar percentages of spatially tuned ROIs across both contexts (Context A: 35.5%, Context: B 39.2%), the MEC FOVs showed slightly more spatial tuning in the second context (Context A: 35.5%, Context B: 42.7%, Figure S6D). Between the EC projections, similar proportions of the spatially tuned axons were significantly tuned in both contexts (32.1% of LEC ROIs and 30.3% of MEC ROIs, Figure S6E). When we investigated these axons that were tuned in both contexts, we found that MEC axons’ place fields tended to move further than those of the LEC axons (mean distance LEC: 47.6 cm, MEC: 85.1 cm, Figure S6E). We also observed that the LEC ROIs showed notable enrichment at locations just prior to the reward zone locations while the distribution of the MEC ROI centroids was much more uniform (Figure S6F). These results are consistent with the reward location biasing reward the activation and tuning of the LEC projection based on the distance to and especially proximal to the reward location. Additionally, that changing the visual VR context is sufficient to induce a remapping of the spatially tuned MEC population, similar to what has been observed previously (Cholvin et al. 2021) and in CA1 pyramidal cells (Aronov & Tank 2014, Priestley et al. 2022).

To directly assess how well the two EC projections encode position information, we attempted to decode both context and location from the activity of EC axons. We found that it was, in fact, possible to predict both variables from either subregion with similar accuracy (context decoding, LEC: 64.5%, MEC: 70.7%, Figure 4F). Both MEC and LEC datasets yielded good position decoding performance when the classifier was also successful at predicting the contexts (average by mouse of median decoding error, LEC: 16.3 cm, MEC: 11.8 cm, *p* = 0.2217 unequal variances t-test); however, when the context was incorrectly decoded, the LEC dataset produced significantly better predictions of the position (relative to the lap start and the reward location) compared to the MEC data (average by mouse of median error, LEC: 35.3 cm, MEC: 72.7 cm, *p* = 0.0007 unequal variances t-test, Figure 4G). We further observed that the position decoding error resulting from the LEC axons was much smaller compared to MEC if we trained exclusively on the opposite context than we decode on (Figure 4H). Lastly, we asked if it would be possible to decode the previous context during the ITI. We find that it is indeed possible to resolve which context the animal’s recent experience from both the activity of both the LEC and MEC axon populations (Figure 4E, S6B). Taken together, these results indicate that the MEC axonal activity in CA1 contains more distinct representations of space between contexts, while the LEC axons are more biased by the reward location.

### Shuffled goal location diminishes spatial selectivity of LEC-to-CA1 representations

Given the aforementioned tendency of goal locations to stabilize spatial activity in the LEC axons, we next tested what would happened if the reward was not in a fixed, predictable location, but instead at a novel location on every lap (Figure 5A, S7B). In this case, we observed, that comparing the activity of even to odd numbered laps, the MEC axonal input to CA1 appeared to be more spatially stable than did the LEC input (Figure 5B). In fact, comparing even to odd laps resulted in higher spatial PV correlations in all MEC recordings as compared to LEC (PV correlations odd/even laps mean, LEC: 0.195, MEC: 0.295, Figure 5C). Additionally, during this shuffled rewards task, a higher percentage of MEC axons exhibited significant spatial tuning (percent of axon ROIs spatially tuned, LEC: 22.2%, MEC: 40.1%, Figure 5C) and decoding the position of the animals was more successful from the MEC axons than from the LEC population (median decoding error LEC: 34.4 cm, MEC: 17.1 cm, Figure 5C). However, we do note that while there were fewer spatially tuned LEC ROIs, we did see some ROIs with significant spatial tuning; centroids of spatially tuned ROIs from both MEC and LEC axons spanned the entire track (Figure S7D). Given that reward is a particularly salient cue, we also checked whether adding additional salient, multi-model spatially-fixed stimuli would increase the spatial properties of the axonal ROIs. Adding both spatially fixed olfactory and auditory cues, we found modest increases in percentages of spatially tuned ROIs as well as improved performance of a decoder at classifying the animals location (Figure S7A-C). We did not observe any specific enrichment of spatially tuned cells in the vicinity of the spatial cues (Figure S7D), suggesting that the reward-driven enrichment we observed in the LEC axons in all previous tasks is unlikely to be purely related to the salience of the stimulus presentation.

**Figure 5.**
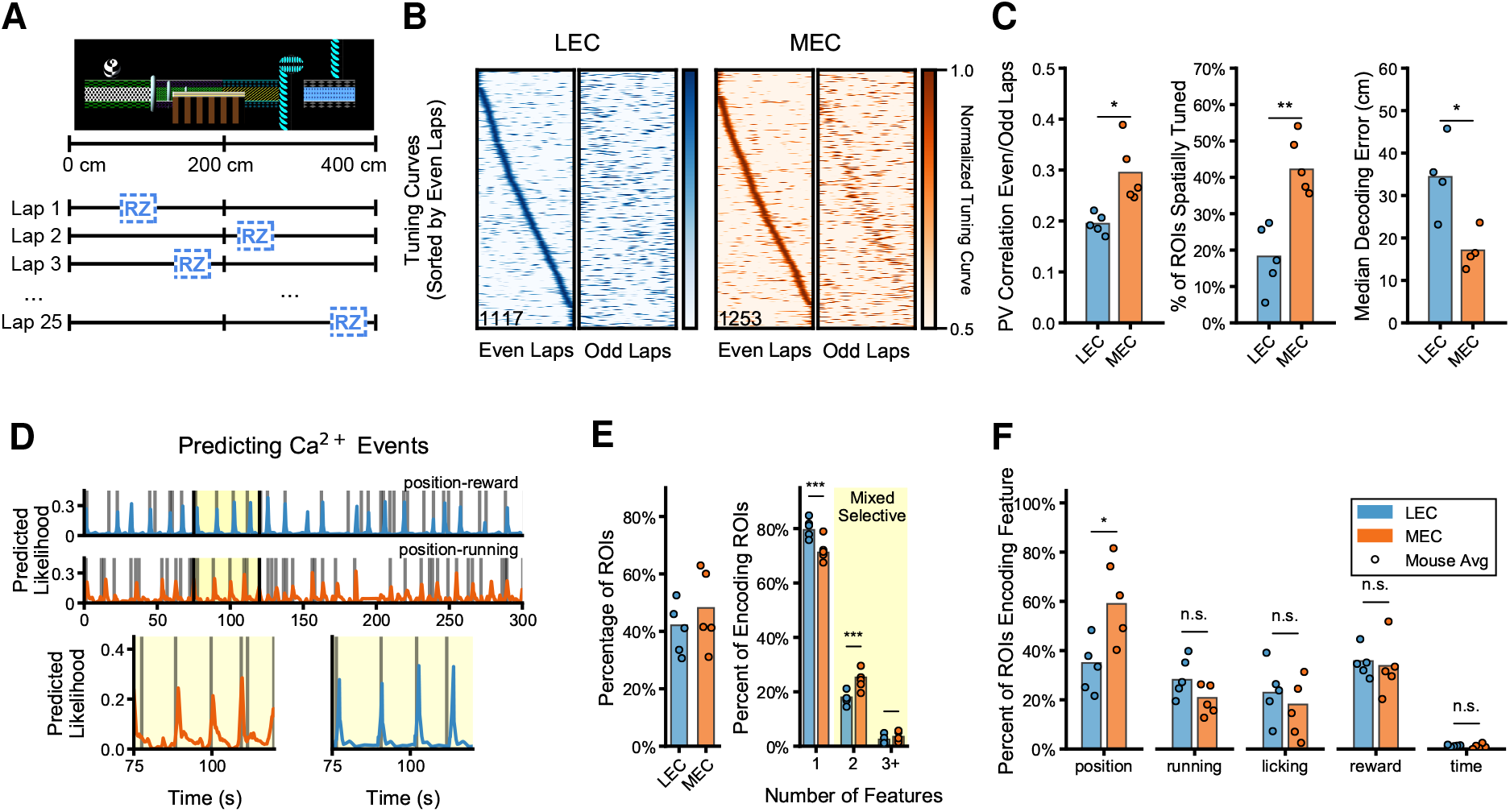
LEC axons showed diminished spatial tuning when reward location was changed every lap. (**A**) Diagram of the shuffled rewards task where the reward location was moved to a random location on every lap. (**B**) Tuning curves for all ROIs in a single FOV of LEC axons (*left*) and MEC axons (*right*) recorded during the randomized reward task. ROIs sorted by the peak Ca^2+^ event rate across all even numbered laps. (**C**) The MEC axons’ activity was significantly more spatially stable then the LEC axons across several measures during the randomized rewards task. *Left*. Spearman’s rank correlation coefficient (ρ) calculated on a population vector of mean activity by position bin for all ROIs between even and odd laps showed higher spatial correlations for MEC axons than LEC. Mean correlation LEC: 0.19, *n* = 5 mice, MEC: 0.29, *n* = 5 mice, Welch’s unequal variances t-test *t* = 3.5301, *p* = 0.0177. *Middle*. More significantly spatially tuned axons (*p* < 0.05 compared to shuffle) were observed in MEC as compared to LEC. Mean across mice LEC: 18.3%, *n* = 5 mice, MEC: 42.2%, *n* = 5 mice, Welch’s unequal variances t-test *t* = 4.8033, *p* = 0.0014. *Right*. Decoding position bin from the activity of MEC axons is significantly more successful than LEC. Median error decoding the animal’s virtual location was calculated for 1000 randomly selected ROIs per mouse, 3 different randomly selected splits were used and results averaged LEC: mean error 34.4 cm, *n* = 5 mice, MEC: 17.1 cm, *n* = 5 mice. Welch’s unequal variances t-test *t* = 3.3423, *p* = 0.0247. (**D**) *Top*. Example performance of predicting calcium transient event times (gray) with modeled transient probability for an LEC (blue) and MEC (orange) ROI. *Bottom*. Expanded highlighted areas from the top traces. (**E**) *Left*. Successful models showed a significant (*p* < 0.5) improvement over a mean firing rate model. Similar numbers of LEC and MEC ROIs were successfully modeled, mean percentage across animals, LEC: 42.1%, MEC 48.1%, Welch’s unequal variances t-test *t* = 0.9058, *p* = 0.3955. *Right*. ROIs best modeled by including more than one type of predictors were determined to be mixed-selective. Mixed selective ROIs can be found in both LEC and MEC. Mean across mice, LEC: 20.5% mixed-selective, *n* = 5 mice, MEC 28.8% mixed-selective, *n* = 5 mice. (**F**) Percent of ROIs with models that include each type of feature. A higher percentage of MEC axons, 60.0% relied on position features than LEC, 34.96%, mean percentage by mouse MEC was significantly higher than LEC, Welch’s unequal variances t-test *n* = 5 mice LEC, *n* = 5 mice MEC, *t* = 3.1429, *p* = 0.0173. Percent of axons was greater than 0 for all types of features, one sided, one sample t-test, Position - LEC: *t* = 3.5093, *p* = 0.0011, MEC: *t* = 4.0250, *p* = 0.0006, Running - LEC: *t* = 4.0032, *p* = 0.0007, MEC: *t* = 3.6192, *p* = 0.0010, Licking - LEC: *t* = 2.2482, *p* = 0.0054, MEC: *t* = 1.4949, *p* = 0.0202, Reward - LEC: *t* = 6.5481, *p* = 0.0001, MEC: *t* = 3.2453, *p* = 0.0014, Time - LEC: *t* = 4.3846, *p* = 0.0005, MEC: *t* = 1.1498.

Since randomizing reward locations resulted in lower percentages of ROIs in the LEC to be spatially tuned, we asked if there were other behavioral factors that could be contributing to the activity measured in the EC axons. We used sets of features related to the animals position, running/velocity, licking, reward presence, as well as the elapsed time (both within a lap as well as since the start of imaging), and found that for some ROIs, these features were able to significantly predict the timing of Ca^2+^ transient activity (Figure 5D, S7F, G). We applied a model search technique (similar to what has been previously described, Hardcastle et al. 2017) in order to find which model best described the responses of each ROI while using the minimal number of features (Figure S7E). Despite the difference in spatial tuning, we were nevertheless able to generate models of the activity in a similar proportion of both LEC and MEC axons, and these models significantly outperformed a mean firing rate model (average per mouse LEC: 42.1%, MEC: 48.1%, difference not significant, *p* = 0.3955 unequal variances t-test, Figure 5E). Further, we found that both LEC and MEC projections contained a significant set of axons that were better described by a mixture of features than by a single type of predictor, similar to previous reports of models generated from recordings from neurons in MEC (Hardcastle et al. 2017) and CA1 (Stefanini et al. 2020) (percent mixed selective LEC: 20.5%, MEC: 28.8%). However, during the foraging task, we did observe a significantly higher percentage of the MEC axons as mixed-selective compared to LEC (Figure 5E). When we looked at which categories of features were most descriptive, we also found that a larger percentage of the MEC axonal population was specifically influenced by the animal’s location compared to LEC, whereas the other features we tested were more evenly represented by each projection (Figure 5F, S7H). Since episodic time coding has previously been described in recordings from the LEC (Tsao et al. 2018), we included a time feature in the model that captures elapsed time from the start of imaging as well since the start of each trial. In our data, the time feature turned out to be unimportant for most of the ROIs recorded. This observation could, however, be due to a limitation of our imaging technique: decoding based on the start times of calcium transients might obscure the ramping activity on a short time scale that occurs on each of the 4m laps. The discrepancy between our findings and the previous study may also lie in the population of cells imaged: while Tsao et al. found ramping activity in all layers of the LEC, they did not specifically target the population of cells that projects to CA1. In this respect, our experimental design may more accurately capture the specific types of representations (multi-faceted but not temporal) present in the LEC projection to hippocampus.

Lastly, we also modeled the activity of the entorhinal input to CA1 as mice performed the alternating rewards and the context alternation tasks (S8). For these models, we explicitly used the distance to the reward as a predictor and separated position labels by context. We found that a higher percentage of axons from either subregion could be successfully modeled during the during the tasks which had fixed reward locations as compared to the random foraging task (Figure S8A). Additionally, when distance to reward was included as a predictor to the model, we no longer saw a significant difference in the proportion of mixed-selective axons in MEC compared to LEC (Figure S8A). Most importantly, we observed that, regardless of the task, there were consistently more MEC (vs LEC) axons described by models that include the position in context, while higher percentages of LEC (vs MEC) axons were better modeled by a distance-to-reward or reward vector type coding of space (Figure S8C).

## Discussion

Our study provides a direct comparison of the LEC and MEC input to CA1 during a series of behaviors that elucidate unique and overlapping roles for each projection. Our comparative approach uncovers both significant spatial tuning and the ability to decode position in both projections. The LEC spatial maps exhibit a robust goal driven enrichment, while the MEC exhibits more within-context spatially uniform representations: together LEC and MEC provide CA1 with both goal-centric and context-centric representations of space, respectively. In 1D tasks, both of these representations could be used to form and stabilize the kind of mnemonic and spatial ‘dual’ map that has been observed in CA1 (Eichenbaum & Cohen 2014, Schiller et al. 2015). Unexpectedly, we find that during goal directed navigation it is possible to decode current location from the LEC input to CA1 (in addition to MEC) at fairly good resolution and throughout the entire 4m context (Figure 1L, S4C). It is possible that neurons downstream of the EC in CA1 might bias representations toward whichever map is the most informative or confident for a given task. These results have important implications for the interpretation of previous studies that manipulated EC inputs to CA1, as well as for our understanding of the various routes spatially informative information takes as it propagates through the cortex to the hippocampus.

LEC has previously been reported to exhibit a low prevalence of spatially tuned cells (Yoganarasimha et al. 2011, Deshmukh & Knierim 2011), which has led to both a series of models (Bush et al. 2014, Rolls et al. 2006, Solstad et al. 2006) and experimental paradigms investigating how MEC activity (and grid cells in particular) might yield place cell activity in CA1 (Hafting et al. 2005, Brun et al. 2008). The current study is unique in that we not only considered MEC inputs, but also characterized the spatial tuning of LEC inputs to CA1 during a variety of tasks. Our results confirm spatial selectivity in MEC-to-CA1 projections, though not necessarily grid-like tuning, but we also find that with the correct task parameters, namely when the animal has a clear navigational goal, the LEC input to CA1 is also spatially stable (Figure 1F, G). We validate this finding by showing that the location of the animal can be decoded from the activity of the LEC as well as the MEC projection to CA1 (Figure 1L). While this observation might seem initially surprising, it is in fact in line with the idea that the LEC may form context, object, location associations as well as vector based object tuning (Deshmukh & Knierim 2011, Tsao et al. 2013, Wilson et al. 2013). Since the reward location would also serve as a prominent proximal landmark in 1D space, our result is also in agreement with models that predict MEC to be involved in more allocentric coding while the LEC maps of space may be considered more egocentric (Lisman 2007, Wang et al. 2018). However, especially in tasks with fixed goal-context pairing, our results indicate that the LEC representations could also be capable of providing the necessary spatially tuned input to drive place field formation, stability, and subsequent remapping with context change. Since location can be decoded from the LEC axonal population throughout the entire 4m track, not just around the goal location (Figure S4C), and since distinct contexts are still represented differently (Figure 4C, D), there is likely some overlap in the tuning profile of the two regions. The potential for overlapping spatial representations between the EC subregions has important implications for studies which investigate cortical input to CA1 through MEC specific lesions and inactivations (Brandon et al. 2014, Hales et al. 2014, Schlesiger et al. 2018), suggesting that both MEC and LEC may work in conjunction to support the tuning and remapping of CA1 place cells. Furthermore, LEC may be providing some spatial and contextual information to CA1, explaining previous reports of animals with lesioned MEC still exhibiting place field formation and remapping (Schlesiger et al. 2018).

When goal locations change, we observe a subsequent reorganization preferentially in LEC-CA1 projection and, importantly, overlapping representations of the goal areas and of areas just prior to goal locations (Figure 2D, S5B, E). Goal related enrichment is also specifically present in the LEC axons between distinct contexts, along with increased PV correlations at locations proximal to the rewarded location (Figure 4D), indicating a possible goal-distance driven representation of space. Using logistic regression to model the Ca^2+^ transients directly, we find specific evidence for goal-distance tuning and that it is significantly more prevalent in the LEC axons compared to MEC (Figure S8C). This result is in agreement with models that suggest that the LEC might be important for CA1 representations of more mnemonic aspects of behavior; however, the representations we observed cannot be accurately described as exclusively “non-spatial”, especially since the activity is sufficient to describe the animals location, even at distances that are far from goal locations (Figure S4C, B). Recent evidence shows dopaminergic reward expectation signals during non-spatial olfactory tasks, mitigated by a pathway between VTA and superficial LEC (Lee et al. 2021). Besides this pathway, the LEC likely receives spatially tuned inputs from postrhinal cortex (LaChance et al. 2019), collateral input from the MEC (Nilssen et al. 2019), or internally from deep layers (via CA1) (van Strien et al. 2009). The existence of these inputs suggests that features of memory and space may be integrated upstream of CA1 within LEC. Whether the dopaminergic projections from the VTA play a role in goal learning during navigation is a question outside the scope of the present study; future studies could help us to further understand how split navigational obligations arise in the EC subregions. Nonetheless, it is notable that the stability of reward-proximal place cells in CA1 have been shown to be uncorrelated with pre- and post-experience activations during offline-memory consolidation-epochs (Grosmark et al. 2021), suggesting that there might be unique mechanisms that contribute to the maintenance of these place fields. An enrichment in stable LEC spatial fields proximal to goals implies that less synaptic plasticity within CA1 is necessary to maintain CA1 place fields in these locations.

It is notable that other studies have found reward-based tuning in both MEC grid and non-grid spatial cells that appears to depend on the task structure (Boccara et al. 2019, Butler et al. 2019). While our results point to a more prominent role for the LEC in goal enrichment during oriented navigation, we do also observe improved position decoding along with increased place field stability in the MEC projection to CA1 around goal locations (Figure 2D, S4C), in agreement with the aforementioned studies. These studies were also targeting grid cells during 2D navigation and did not directly sort cells by whether they participated in the direct projection to CA1. Moreover, models of place field formation predict that CA1 dendritic plateau potentials are necessary for place field formation events, but may not be required for the *maintenance* of place fields (Grienberger & Magee 2022). The uniform distribution of spatial tuning we observe in the MEC input to CA1 combined with an uneven sampling of space, in fact, would be predicted to contribute to an over-representation of reward-adjacent place fields (Dupret et al. 2010). Therefore, the contribution of the goal anticipation enrichment observed in the LEC axons may not exclusively be responsible for the previously observed congregation of CA1 place cells around goal locations (Dupret et al. 2010, Zaremba et al. 2017, Gauthier & Tank 2018) and MEC axons likely have a complementary role. It is also possible that different task dependent mechanisms may contribute to place field formation and enrichment and that the spatial maps observed in CA1 reflect conjunctive processing of both EC inputs, which are unlikely to be fully orthogonal in their representations of space. In fact, there is a gradient of the LEC/MEC projection synapses onto CA1 pyramidal cells which changes along the transverse (Masurkar et al. 2017) and radial axis of the hippocampus (Soltesz & Losonczy 2018), so it is possible that the CA1 response to EC inputs might preferentially weight the LEC vs MEC based on where the postsynaptic cell is located.

As expected, we do find significant spatial information in many of the MEC-originating axons we recorded (Figure 1I). Further, we find that this activity remaps with global changes to the navigational context (Figure 4B, C, D), as has been previously published (Cholvin et al. 2021). Unlike LEC, the stability of the MEC input to CA1 is a fairly reliable predictor of the animal’s location following manipulations to goal locations, which we demonstrate by decoding location across reward zone shifts, with reasonable success (Figure 2F). It has been suggested that grid cell activity in this MEC projection might be fundamental to the development of place field tuning in CA1 (Samsonovich & McNaughton 1997, Bonnevie et al. 2013, Burgess et al. 2007, McNaughton et al. 2006, Fyhn et al. 2002); however, we find no specific evidence that grid-like tuning is over-represented in the observed MEC layer III projection (Supplementary Materials, S9)–in agreement with the previous study (Cholvin et al. 2021). For putative grid-like axons, as nominated by standard methods of classification, we took additional steps to resolve best fit parameters for the potential underlying 2D grid spacing and trajectory (Yoon et al. 2016). We find the LEC and MEC data to “fit” equally well to grid models, indicating that at the population level, the MEC activity is no more “grid-like” than we observe in any multi-peaked spatial cells constrained by the parameters of the 1D navigation task (with a 4 m 1D environment). Most importantly, the grid cell axons would be expected to represent the same size grids (preserved λ) despite remapping trajectories between contexts (Hafting et al. 2005); after the context change, we observe no continuity in the parameters that best fit grid activity to the recorded activity. While it is possible that our result is limited by the length of the track and the resolution of the GCaMP6s indicator, we observe no bias in our recorded MEC populations towards grid-like tuning, which would be expected if the majority of this input was indeed following those kinetics. We specifically see evidence for spatial, likely non-grid, cells with context dependant tuning curves participating the MEC projection to CA1.

In addition to representing the current location and experience, activity in CA1 has also been reported to encode information about the recent past, or “latent information” (Keinath et al. 2020). In our experiments, we see encoding of this kind of memory about recent experience in the cortical inputs to CA1. The ability to accurately decode the previous reward condition (especially in LEC, Figure 2G), as well as the previous context (Figure 4E) during the 2 second timeout between laps, when there are no visual cues, indicates that there is some persistence in the state of the EC neuronal population that is not dependant on the current sensory experience. This possibility is congruent with models of EC circuits following attractor-like dynamics (Yoon et al. 2013). We also observe, mainly in the LEC, a representation of the current navigational goal (Figure 2H). In the present study we cannot conclusively discern whether this finding reflects a retrospective coding of the recent past or a prospective coding for the animal’s planned future action on the next lap. In fact, the EC has previously been reported to maintain a memory of object manipulations, with cells tuned to the location of novel or displaced objects (Deshmukh & Knierim 2011, Tsao et al. 2013). Additionally, goal related cells found the human EC have been shown to reactivate during hold/pre-trial periods (Qasim et al. 2019). This observed object trace activity could be indicative of remembering important or changed locations, which could be of future interest and could play an important role in future route planning.

Taken together, our results suggest distinct, yet overlapping roles for LEC and MEC in crafting the spatial and mnemonic maps observed in CA1; thus, we build upon recent observations which challenge the traditional strict dichotomy (Nilssen et al. 2019, Save & Sargolini 2017). Our study has the benefit of specifically probing the input to CA1 through axonal imaging and examining how the intact circuit performs–our unique combination of this technique and perspective allowed us to conduct a direct comparison between the LEC and MEC axonal activity. Of course, several questions remain: for instance, an investigation into how reward expectation signals accumulate in the LEC could be a potential direction for future inquiry, however, it is reasonable to speculate that CA1 partially inherits goal-skewed spatial tuning directly from the LEC given our direct observation of these signals. Finally, our observations imply that the LEC likely contributes to spatial activity in the absence MEC input. While outside the scope of the reported experiments, additional differentiation between the subregions may become apparent with increasing task complexity.

## Methods

### Mice

All experiments were conducted in accordance with the NIH guidelines and with the approval of the Columbia University Institutional Animal Care and Use Committee. All animals used for experiments were 6-12 week old, male C57/B6J mice. Animals were kept on a 12 hour light/dark cycle and housed in the vivarium at Columbia University. Animals were kept in groups of 2-5 with litter mates and had *ad libitum* access to food. During training and imaging, mice were water-restricted and maintained at least 80% of their pre-restriction weights as determined by daily weighing of mice. Well trained animals received the majority of their water during experiments, however 2-10 minutes of supplemental free access to water was given to animals, as necessary to maintain their weight above the 80% threshold.

### Surgery

#### Injections

Mice were stereotactically injected in either the LEC or MEC with a mixture of 2/3 rAAV1-*Synapsin*-GCaMP6s and 1/3 mixture of rAAV9-*hSynapsin*-DIO-tdTomato and h56d-Cre (LEC: relative to bregma −3.0 mm AP, −4.7 mm ML, −2.6 mm, −2.5 mm, −2.4 mm DV, MEC: 0.2 mm rostral of the transverse sinus, - 4.7 mm ML relative to bregma, −1.3 mm, −1.2 mm, −1.1 mm DV). 2 pulses of 28.8 nL were injected at each DV location (depth) using a Nanoject syringe. Additionally, for all animals, 50x diluted *CamKII*-NLS-tdTomato was injected into CA1 to sparsely label the pyramidal cell layer (−2.3 mm AP, −1.5 mm ML, −1.1 mm, −1.0 mm, −0.9 mm DV relative to bregma). 2 pulses of 32.2 nL were applied at each DV target in CA1.

#### Window Implants

Following the viral injections, mice were give 3 days to recover. Then, a craniotomy was performed directly over left dorsal CA1, and a 3 mm imaging window and cannula were implanted in accordance with the previously described procedures (Kaifosh et al. 2013, Lovett-Barron et al. 2014). Custom 3D printed titanium head fixation bars were then attached to the skull with dental acrylic.

### Behavior

After recovering from the craniotomy, mice were habituated to head fixation, then water deprived and trained to lick at a water spout while running on a wheel. Movement of the running wheel advanced the position of visual VR cues, which were displayed on 5 LCD monitors surrounding the mouse. The running wheel and VR setup were as previously described (Warren et al. 2021, Priestley et al. 2022). Mice were trained first to lick for water rewards spread across the entire track, and then, eventually, for a single fixed reward zone. Reward zones were fixed 20 cm sections of the track. When mice entered the reward zones, they were given an initial drop of water followed by a time period (”reward duration”) during which animals could lick at the reward spout and receive additional water droplets–provided they stayed within the 20 cm area. Mice that did not slow down would run past the rewarded area and not be able to get additional reward; thus the animals had an incentive to learn the reward locations and not fully rely on the initial drop as a cue. Reward durations were initially 10 - 15 seconds and then reduced as animals learned the task. Animals were considered proficient to start experiments after they would run for at least 50 laps in 20 minutes while reward duration was 2 seconds. The training VR context was distinct from those used during experiments; however, for the data shown, all animals had at least 1 prior experience in the environments. All VR environments used during the experiments were 400 cm long with a start and end “tunnel” at each side that the animal would exit from and enter into at the beginning/end of each lap. Following each lap, there was a 2 second timeout when all of the visual cues were dark prior to the next lap starting.

#### VR Navigation

Mice first had to traverse 25 laps in a VR Off context without the visual cues: the only spatial cue was the reward zone, which was fixed at a location 360 cm after the start of the lap. Once mice reached 400 cm, there was a 2 second timeout before the next lap started. After 25 laps of running without visual cues, the animals completed 25 laps with access to the visual stimuli–the VR On context. The reward zone remained fixed in the same location and the 2 second timeout did not change. After the VR On laps, mice would continue to run in a subsequent VR Off block of laps; however, data from this block was not included in our analysis (other than the lap-by-lap correlation plot, Figure 1G).

#### Reward Alternation

For the reward learning task, we used two reward zones, RZ1 at 360 cm and RZ2 at 110 cm relative to the start of the virtual track. The reward location was fixed at RZ1 for a block of 10 laps, then moved to RZ2 for 10 laps, then returned to RZ1 for 10 laps, and finally moved once again to RZ2 for 10 laps. As described above, upon entry into the reward zone, an initial drop of water cued the animals to the reward location; additional rewards were available if the mouse learned to stop and lick within 20 cm of the reward start for a period of 2 seconds. There was no visual cue that changed or indicated which reward zone was currently active.

#### Context Alternation

To test the EC axons during complete contextual shifts, we used two completely distinct sets of visual cues along the 400 cm track. Mice were first presented with a Context A for 10 laps, and then Context B for 10 laps, followed by Context A and then B each for another 10 laps. The reward location remained fixed at 360 cm for both contexts. Context A was the same context used for the VR navigation and reward alternation tasks, so the mice may have been more familiar with it; however, for the data presented here, mice had also run at least 20 laps in Context B on a previous day, suggesting some level of familiarity with both contexts. We did not explicitly test for any effect of novelty during the context alternation task.

#### Random Foraging

For the foraging task, we used a distinct fixed VR environment that the mice had never previously experienced. This new environment contained a single reward location that was moved to a new position on every lap. The location of the reward was still cued on every lap with an initial drop of water, but mice that responded quickly enough and stopped within 20 cm of the initial drop could consume additional water rewards for 2 seconds. For a subset of mice, after 25 laps of the random foraging task, a multi-sensory manipulation was conducted: a 5 kHz tone was presented at location 265 cm for 20 cm, and an odor appeared at 50 cm for 10 cm. For the tone, a small speaker was positioned near the base of the running wheel. For the odor, a custom 3D printed nose-cone was positioned just in front of the mouse, with a constant airflow moving through a container of either a neutral background (aeorsolized mineral oil) or an odor cue (monomolecular carvone dissolved in mineral oil). Odorants were evacuated from the nose-cone with a vacuum line.

### Imaging

Two-photon imaging was conducted at 30 Hz at 512 × 512 pixel resolution using an 8 kHz resonant scanner (Bruker) with a 920nm laser (Coherent) set to 50-125 mW as measured at the objective. Laser power and alignment were verified each day before starting the experiment. All datasets were collected using a 16X Nikon water immersion objective (0.8 NA, 3.0 mm WD) at 3.5 times digital zoom, resulting in a FOV of 240 *μ*m x 240 *μ*m. Green (GCaMP6s) and red (tdTomato) fluorescence were collected through a filter cube set (green HQ525/70 m-2p, red HQ607/45 m-2p; 575dcxr, Chroma Technology) coupled to a GaAsp PMTs (Hamamatsu Model 7422P-40) and amplified using a custom dual stage preamp (1.4 × 105 dB, Bruker).

### Preprocessing

#### Motion Correction

Datasets were processed using the SIMA python package for handling sequential imaging data (Kaifosh et al. 2014). Motion correction was accomplished using a custom implementation of the NoRMCorre algorithm (Pnevmatikakis & Giovannucci 2017) which was modified in order to drop low-correlation patches or entire frames which were corrupted by z-motion, as identified by outliers during the template correlations step. In some cases, we also report frame stability, as measured after the motion correction and calculated as the individual frame correlation to the time average of the entire dataset as in Grosmark et al..

#### Segmentation and Signal Extraction

ROIs detection, neuropil subtraction, and signal extraction were achieved used the python implementation of the Suite2p (Pachitariu et al. 2017) package. The visual interface was then used to inspect all of the ROIs to ensure there were no spurious components detected. Axon ROIs were taken to be either individual boutons or small segments of axons.

#### Δ*F/F and Correlations calculation*

The fluorescence was smoothed with a Savitzky–Golay filter (window length 51, polynomial order 3) and then a baseline was calculated using a max min filter (window size 500, *σ* 10); finally, Δ*F/F* was calculated. In order to ensure that axons were not over-segmented, Pearsons’s pairwise correlation coefficient was calculated between the Δ*F/F* traces for all pairs of ROIs detected in a FOV. ROI pairs with an extremely high correlation coefficient (> 0.7) were considered to be from the same axon segment and merged.

#### Transient Detection

Transients were detected and used for all subsequent analysis unless specifically annotated as otherwise. Transient detection was done with a threshold of height based on an estimate of the noise as the standard deviation of the signal. Transients started when the Δ F/F surpassed 2*σ*, and were completed once the signal dropped below *sigma*. To be counted as a transient, the signal had to pass 2.5*σ* and last for at least 1 s. Within-transient times were considered unobserved for the purposes of the subsequent analysis. Iteratively, the transients were removed and *σ* was re-estimated in order to reduce any skewing effect of with-transient times, similar to previously used approaches (Danielson et al. 2016). Axons with very few detected transients (< 0.25/min) or very poor quality signals (median within transient maximum fluorescence < 3.25*σ* or 0.35Δ*F/F*) were not included in the analysis.

### Analysis

#### Spatial Tuning

In order to identify axons as spatially tuned, we calculated the spatial information (Skaggs & McNaughton 1993) based on the Ca^2+^ transient onset times for each ROI. Then we circularly shuffled the position of the transients on each lap and recomputed the spatial information. Axons that had spatial information scores higher than 95% of shuffles (*p* < 0.05) were considered significantly spatially tuned. Significance of spatial tuning was calculated independently for the conditions considered (VR On/VR Off, RZ1/RZ2 Context A/Context B, and for the random foraging task Visual Only/Multi-Sensory).

#### Place field detection

Place field locations for each of the spatially tuned axonal ROIs were calculated using a method similar to those previously reported for CA1 pyramidal cells (Priestley et al. 2022), except using the transient event onset times we recorded from the EC axons. Briefly, we identified where in the virtual environment axons’ transient onsets occurred more often than expected by chance. In order to do this, we calculated the occupancy-normalized tuning curve for each axon, for epochs in which the animals were actively navigating through the virtual environment (velocity > 5 cm/s). Then, we calculated 1000 turning curves based on within-lap circular shuffles of the position. Positions where tuning curves that exceeded 95% of the shuffled data for at least 4 cm were considered candidates for place field locations. The place field center and width were assigned by fitting a Gaussian to the position of the animal at each of the transient Ca^2+^ events (center as the mean of the distribution and width as 2*σ*). Place fields were additionally required to have at least 3 Ca^2+^ transients within them (to filter out low activity axons). Place fields were calculated independently for each of the conditions considered (VR On/VR Off, RZ1/RZ2 Context A/Context B, and for the random foraging task Visual Only/Multi-Sensory).

#### Position Decoding

In order to make comparisons about the degree to which EC input to CA1 might be used for the propagation of spatially selective signals, we trained a multi-class (one-vs-one) Support Vector Machine (as in Stefanini et al.) on the extracted transients. For each dataset, a random sample of ROIs was used to ensure that the same number of ROIs were used per mouse for both LEC and MEC animals. For all position decoding analyses, the track was split into 4 cm bins each given a separate class label. Two laps were then held out of the dataset and the classifier was tested on these 2 while being trained on the rest. The train/test split was rotated across all laps until each lap was tested on once. ROIs used for decoding as well as train/test splits were determined psuedo-randomly for 3 different splits. Unless annotated, 1,000 ROIs were used per FOV for the VR navigation and random foraging tasks, 900 ROIs were used for the reward alternation task, and 750 for the context change tasks. FOVs without sufficient ROIs detected were excluded from this analysis. For the VR navigation task, each condition (VR Off or VR On) was considered separately, and only laps 5-25 for each condition were analyzed. For the reward alternation task, RZ1 and RZ2 conditions were also considered separately; meanwhile, for the context alternation task, positions in Context A and Context B were given different class labels, and all 40 laps were included in a single analysis (expect for the across context decoding). For all of the position decoding analysis, locations 0 - 25 cm and 375 - 400 cm were not included due to most of the visual cues being blocked by the end tunnel and for the randomized reward task, the first lap was excluded from the analysis (to make an even number for the train/test split).

#### Velocity Decoding

To decode the speed of the mice from the Ca^2+^ activity, we used Lasso Regression as in (Stefanini et al. 2020), training on the Δ*F/F* from a randomly selected subset of 1000 axonal ROIs per mouse. Leave one out cross validation by lap was used to asses model performance.

#### Context/Reward zone Decoding

Previous lap context or reward zone were decoded similarly during the 2 second between-lap timeout period when the VR screens were dark. Decoding was performed by training a linear SVM on a smoothed (Gaussian smoothing *σ* = 0.27 s) vector of the Ca^2+^ transient onset times, specifically from the inter-lap period. The classifier was trained on 36 laps and tested on 4, one from each context/RZ block (test laps were the identical lap-in-context relative to the last time the context switched).

#### Grid Classifier

The grid tuning of axons was assessed as a two-step process. The first step was to find identify putative “grid-like” ROIs with activity that followed a spatially localized periodic firing pattern; the second step was to fit those ROIs to an idealized grid pattern, as in Yoon et al.. The identification in the first step was based on a series of criteria adapted from Domnisoru et al. and similar to commonly used, established criteria used to classify MEC activity as “grid-like” in 1D environments (Yoon et al. 2016, Gu et al. 2018, Cholvin et al. 2021). Specifically, axons had to meet 5 criteria: (1) there must be 2 identified spatial fields on the track, (2) the number of transitions between in-spatial field and out-of-field firing must be L/ (5*w*) where *L* is the track length (in this case 400 cm) and *w* is the mean spatial field width of the axon’s 1D response, (3) the width of the widest spatial field for any axon must be less then 5*w*, (4) At least 30% of the spatial bins must be able to be assigned to either in-field or out-of-field periods, (5) the observed in-field activity must be twice that of out-of-field activity. Given that a lot of presumably non-grid LEC axons also passed these criteria, we took the additional step to fit the 1D tuning curves to the best matching idealized tuning curve by taking a linear slice through a 2D grid lattice. As developed in Yoon et al., this problem equates to resolving the origin **c** = (*c*_1_, *c*_2_), period, λ, and trajectory, or angle relative to the x-axis, *θ*, in a 2D coordinate space defined by infinitely repeating equilateral triangles. The ideal grid firing rate was approximated as the sum of three 2D sine waves:

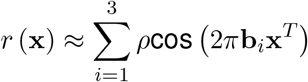

Where 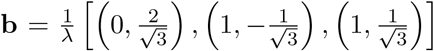, and we set *ρ* = 1 and scale *r* (**x**) accordingly. This was accomplished by, first, finding the spectral peaks of the Fourier transform, *f*_1_ (λ, *θ*) < *f*_2_ (λ, *θ*) < *f*_3_ (λ, *θ*), then solving for 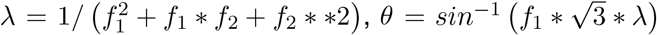 and finally, finding the phase offset using the phase offset of the peaks in Fourier space; with *δ*_1_ = *ϕ* (*f*_1_) and *δ*_2_ = *ϕ* (*f*_2_), then 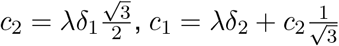. Finally, since fit quality of the analytical solution is limited by finite size effects, we refine the estimate of all parameters through a gradient descent numerical optimization.

#### Logistic Regression Model

In order to determine the relationship between Ca^2+^ transient activity and both the animal’s behavioral state and location in the VR, we used a logistic regression model to predict transient event probability times. With *m* individual features, the event probability, 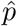 is then:

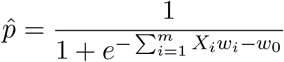

We performed the analysis on groups of features related to position, running, licking, reward, and time such that: **Position** - the 400 cm track was split into 100 position bins, which were encoded using a one-hot technique, so these variables were 1 on time samples when the animal occupied each bin, and 0 otherwise; **Running** - both the velocity of the animals as well as binary features encoding run start, run stop, and running vs non-running frames; **Licking** - binary, set to 1 only on time samples when a lick event was detected; **Reward** - binary vector representing times when the reward is available to the animal, also separate features for reward start and stopping events; **Time** - time was encoded with separate features encoding lap time as well as experiment time (similar to Tsao et al. but modified for the linear VR track) with an additional predictor for the current lap count since the start of the experiment. For all regression modeling, we used 20 ms time bins, and features were smoothed with a guassian kernel (*σ* = 0.8 ms). Lags from 0 to 1.6 s were added to the running features, and ±1.6 s were applied to the licking features. We trained the models using leave-one-out cross validation. Model performance was evaluated based on the log-likelihood of the held out laps compared to a mean event rate model. Just as in Hardcastle et al., axons for which no model out-performed the mean firing rate were considered unclassified. Axons which fit to a single group of features were then trained on double models. Then, after finding the best performing double feature model, we assessed if it performed significantly better than the less complex, single feature model. Axons that were better fit with double feature models were then fit to a triple variable model–this process was repeated until we fit the 5-variable model. Significance was assessed through a one-sided signed t-test with a significance value of *p* = 0.05, using each model trained as part of the cross validation folds as a sample. For the reward and context alternation tasks, we added an additional feature which was constructed similarly to the position features, except that bins were labeled by distance to reward, not relative to the start of the track. Also, for the context alternation task, positions in Context A and Context B were assigned different variables.

## Lead Contact and Materials Availability

Further information and requests for resources and reagents should be directed to the Lead Contact Attila Losonczy. All unique resources generated in this study are available from the Lead Contact with a completed Materials Transfer Agreement.

## Acknowledgments

J.C.B. is supported by NIMH F31NS110316. A.L. is supported by NIMH 1R01MH124867 and 1R01MH124047; and National Institute of Neurological Disorders and Stroke (NINDS) 1U19NS104590 1U01NS115530, and 1R01NS121106. We thank Drs. S. Fusi, J. Jacobs, and R. Costa for comments and discussion on the project and the manuscript. We thank Drs. J. Priestley, Z. Liao, A. Grosmark, J. O’Hare, and all other members of the Losonczy lab for comments and discussion on the manuscript.

## Author Contributions

J.C.B. and A.L. conceived the study. J.C.B. performed experiments and analysed data. J.C.B. and A.L wrote the paper.

## Supplementary Materials

**Figure S1.**
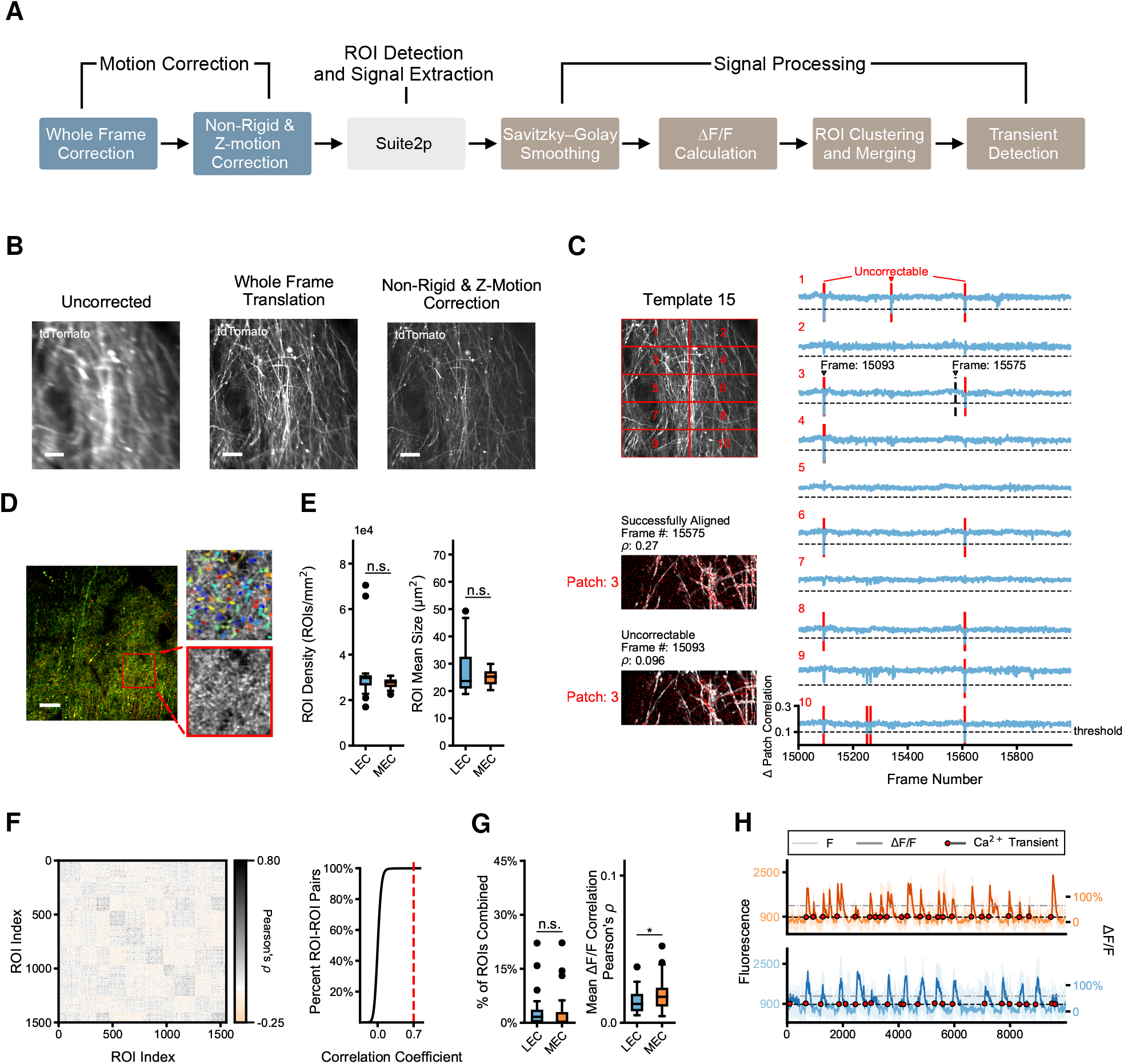
Details of image and signals processing. (**A**) Steps used when processing data collected during all experiments. (**B**) Example results of motion correction algorithm conducted on static tdTomato signal. First a whole frame alignment is performed, then a non-rigid correction step which also identifies and removes uncorrectable patches of data (likely due to z-motion). (**C**) The non-rigid correction template is updated every 1000 frames. *Upper Left*. The data is broken into 8-12 patches which are corrected individually to allow for non-rigid distortions of the data to be corrected. *Lower Left*. Example of single imaging frame data (red) overlayed on the template at shifts corresponding to the peak 2D cross-correlation. Some data cannot be aligned well to the template and the resulting best correlations are low (i.e. Frame #: 15093). *Right*. Outliers in the per patch correlations are identified and removed (red lines). Following the patch alignment, shifts are upsampled to match every row in each patch to ensure smooth boundaries between the patches. (**D**) *Left*. Example time averaged FOV for imaging LEC axons through intact hippocampus. Green - GCaMP6s Red - tdTomato. Scale bar is 25*μ*m. *Right*. Magnified view of indicated area identifying the axonal segments and boutons used to extract signals. Both time average (*bottom*) and overlayed *suite2p* spatial masks (top) are shown. See Figure 1B for example of MEC imaging FOV. (**E**) Details of the ROI identification step. *Left*. No significant difference was found in the mean ROIs extracted per mm^2^ between all LEC and MEC datasets. Welch’s unequal variances t-test, LEC: 2.97*e*4 ROIs/mm^2^, *n* = 32 datasets, MEC: 2.78*e*4 ROIs/mm^2^, *n* = 25 datasets. *t* = 1.5789, *p* = 0.1236. *Right*. Additionally, there was no significant difference in the mean ROI size, LEC: 23.6 *μ*m^2^, MEC: 25.3 *μ*m^2^, Welch’s unequal variances t-test, *t* = 1.6429, *p* = 0.1085. (**F**) Δ*F/F* between pairs of ROIs. *Left*. ROI-ROI correlation matrix for one example axon imaging dataset. *Right*. Person’s *ρ* for all pairs of ROIs from the example show, A small number of ROI-ROI pairs exhibited strongly correlated signals, likely the result of over segmentation of axons branching just outside the FOV. Axon pairs with with high correlations were merged, threshold for merging ROIs indicated by red line. (**G**) Both LEC and MEC datasets showed low correlations between Δ*F/F* between ROIs. *Left*. Percentage of ROIs merged per dataset was not different between LEC and MEC datasets. LEC: 3.1% ROIs merged, *n* = 32 datasets, MEC: 3.0% ROIs merged, *n* = 25 datasets, Welch’s unequal variances t-test, *t* = 0.0843, *p* = 0.9331. *Right*. While correlations between pairs of ROIs was similar was similarly small between the LEC and MEC datasets, the mean correlation coefficient was slightly higher on average for the MEC datasets, LEC: 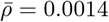, MEC: 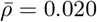, Welch’s unequal variances t-test *t* = 2.0606, *p* = 0.0462 (**H**) Raw fluorescence overlayed with the Δ*F/F* and detected Ca^2^+ transient events. Event detection threshold is initially set at 2*σ* (gray line) and then recalculated with putative event times removed (See *Methods* for details of event detection).

**Figure S2.**
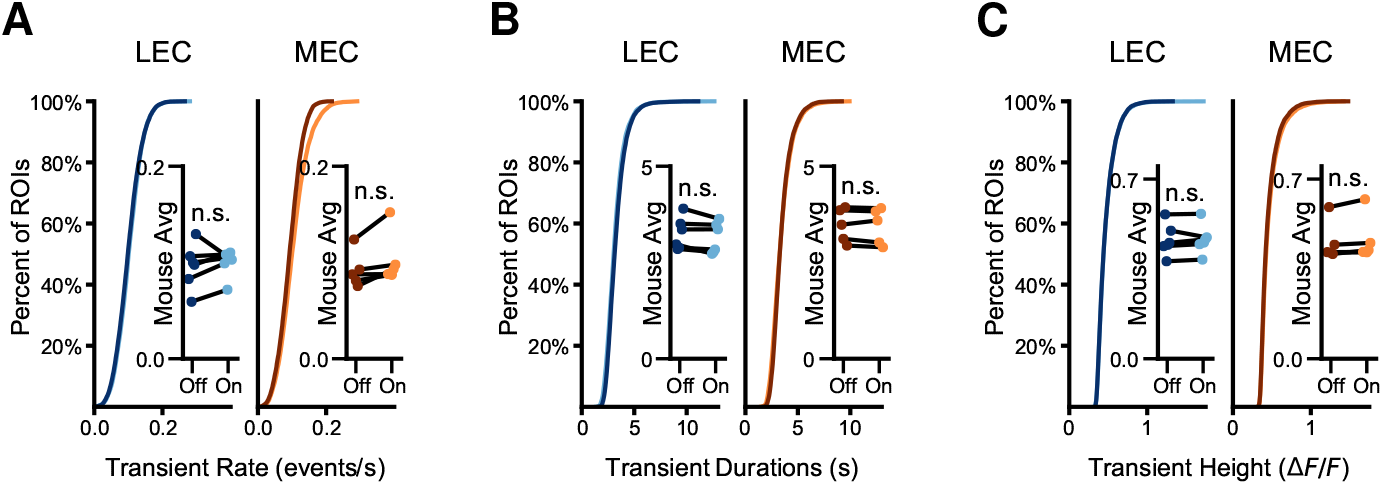
Characterization of GCaMP6s Ca^2+^ Transient Events recorded in EC axonal input to CA1. (**A**) Ca^2+^ event rate was similar for both the VR On and VR Off conditions, but slightly elevated for the VR On condition. LEC: VR Off - 0.1000 events/s, VR On - 0.1022 events/s, MEC: VR Off - 0.0929 events/s, VR On - 0.1041 events/s. Mean event rate per ROI, Two sample Kolmogorov-Smirnov test, LEC: *n* = 7715 ROIs, *D* = 0.0363, *p* = 7.71*e* – 5, MEC: *n* = 6342 ROIs, *D* = 0.1099, *p* = 9.303*e* – 34. *Inset*. Mean rate, by mouse, Two-sided paired t-test, LEC: *n* = 6 mice, *t* = 0.5614, *p* = 0.5987, MEC: *n* = 5 mice, *t* = 2.2668, *p* = 0.0860. There was no significant difference in the by mouse average event rate between the LEC and MEC axons Welch’s unequal variances t-test, VR Off: *t* = 0.2946, *p* = 0.7750, VR On: *t* = 0.3024, *p* = 0.7730. Two-factor ANOVA, effect of VR condition (On vs Off), *F* = 0.6209, *p* = 0.4409, axon type *F* = 0.0003, *p* = 0.9857, interaction of type and condition, *F* = 0.1851, *p* = 0.6721. (**B**) Ca^2+^ event duration was similar in both conditions, but slightly shorter on the laps when VR cues were visible LEC: VR Off - 3.29 s, VR On - 3.18 s, MEC: VR Off - 3.45 s, VR On - 3.43 s. Mean duration by ROI, Two sample Kolmogorov-Smirnov test, LEC: *n* = 7715 ROIs, *D* = 0.0564, *p* = 4.393*e* – 11, MEC: *n* = 6342 ROIs, *D* = 0.0547, *p* = 1.127*e* – 8. *Inset*. Mean duration by mouse, two-sided paired t-test LEC: *n* = 6 mice, *t* = 2.3416, *p* = 0.0662, MEC: *n* = 5 mice, *t* = 0.4819, *p* = 0.6550. There was no significant difference in by mouse mean event duration between the LEC and MEC axons Welch’s unequal variances t-test, VR Off: *t* = 0.8466, *p* = 0.4206, VR On: *t* = 1.1828, *p* = 0.2708. Two-factor ANOVA, effect of VR condition (On vs Off), *F* = 0.1351, *p* = 0.7175, axon type *F* = 0.1080, *p* = 0.1637, interaction of type and condition, *F* = 0.0645, *p* = 0.8023. (**C**) Maximum fluorescence during Ca^2+^ transient events was similar and slightly larger in the VR On condition. Fluorescence values, LEC: VR Off - 0.4712, VR On - 0.4726, MEC: VR Off - 0.4518, VR On - 0.4673, Median height by ROI, Two sample Kolmogorov-Smirnov test, LEC: *n* = 7715 ROIs, *D* = 0.0121, *p* = 0.6293, MEC: *n* = 6342 ROIs, *D* = 0.0374, *p* = 0.0003. *Inset*. Mean by mouse. Two-sided paired t-test, LEC: *n* = 6 mice, *t* = 0.7448, *p* = 0.1129, MEC: *n* = 5 mice, *t* = 2.0249, *p* = 0.1129. There was no significant difference in by mouse mean event height between the LEC and MEC axons Welch’s unequal variances t-test, VR Off: *t* = 0.1934, *p* = 0.8517, VR On: *t* = 0.0154, *p* = 0.9882. Two-factor ANOVA, effect of VR condition (On vs Off), *F* = 0.0413, *p* = 0.8412, axon type *F* = 0.0155, *p* = 0.9022, interaction of type and condition, *F* = 0.0218, *p* = 0.8841.

**Figure S3.**
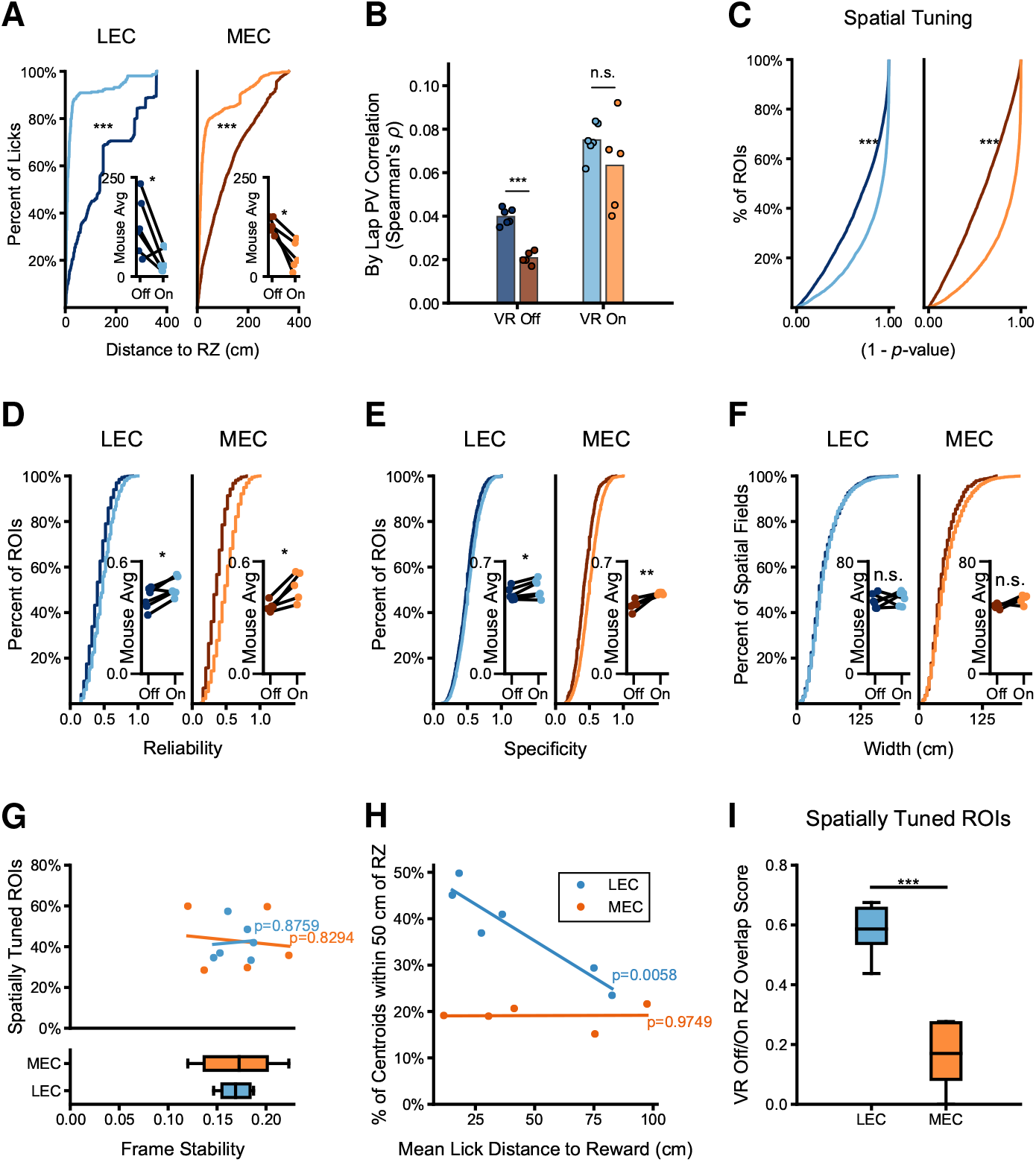
Properties of EC axonal input to CA1 during goal directed spatial navigation task. (**A**) Animals tended to show anticipatory licking at the reward spout when they expected that the RZ was near. Both LEC and MEC labeled mice licked significantly closer to the RZ when the visual cues were turned on as shown by the distance from each location to the RZ, excluding licks which occurred within the RZ. Two sample Kolmorgorov-Smirnov test LEC: VR Off - *n* = 397 licks, mean distance 146.67 cm, VR On - *n* = 308, mean distance 32.17 cm, *D* = 0.6596, *p* < 0.001, MEC: VR Off - *n* = 816 licks, mean distance 124.93 cm, VR On - *n* = 1618 licks, mean distance 46.72 cm, *D* = 0.5200, *p* < 0.001, for mean mouse licking behavior, two-sided paired t-test, LEC: *n* = 6 mice, *t* = 3.0069, *p* = 0.0299, MEC: *n* = 5 mice, *t* = 4.4120, *p* = 0.0116. (**B**) When there were no VR cues, the lap-to-lap PV tuning curve correlations where higher in the LEC axons than in the MEC axons, average Spearman’s *ρ* between PV tuning curve, by mouse, LEC: *n* = 6 mice, MEC: *n* = 5 mice, Welch’s unequal variances t-test *t* = 9.4985, *p* = 5.618*e* – 6. However, during the VR On condition there was no significant difference between the lap-to-lap correlations between the LEC and MEC axons, Welch’s unequal variances t-test *t* = 1.1538, *p* = 0.3021. two-factor ANOVA, effect of axon type, *F* = 10.2752, *p* = 0.00473, effect of VR condition (On vs Off) *F* = 66.4726, *p* = 1.869*e* – 7, interaction of axon type and VR condition *F* = 0.6294, *p* = 0.4379. (**C**) Spatial tuning score for all ROIs recorded as 1 – *p*-value. *p*-value determined based on comparing Skagg’s spatial information score from actual data to Ca^2^+ transient times shuffled on each lap. Spatial tuning with VR cues turned on are significantly higher then for laps when the VR was turned off *Left*: LEC. *Right*: MEC. Two-sample Kolmogorov-Smirnov test LEC: *n* = 7715 ROIs, *D* = 0.2294, *p* < 0.001, MEC: *n* = 6342, *D* = 0.3792, *p* < 0.001. (**D**) Spatial tuning reliability was higher when the mice had access to the visual VR cues. Kologorov-Smirnov test LEC: VR Off - *n* = 1603 spatial ROIs, VR On - *n* = 3319 spatial ROIs, *D* = 0.1608, *p* = 1.041*e* – 24, MEC: VR Off - *n* = 624 spatial ROIs, VR On - *n* = 2782 spatial ROIs, *D* = 0.3403, *p* = 1.118*e* – 51. *Figure inset*. Mean by mouse. Two-sided paired t-test LEC: *n* = 6 mice, *t* = 3.4873, *p* = 0.0175, MEC: *n* = 5 mice, *t* = 3.8653, *p* = 0.0181. There was no significant difference in reliability between the LEC and MEC axons Welch’s unequal variances t-test, VR Off: *t* = 1.5253, *p* = 0.1692, VR On: *t* = 0.1410, *p* = 0.8920. Two-factor ANOVA, effect of VR condition (On vs Off), *F* = 11.0678, *p* = 0.0038, axon type *F* = 0.5346, *p* = 0.4741, interaction of type and condition, *F* = 0.9350, *p* = 0.3464. (**E**) Spatial tuning specificity was higher when the mice had access to the visual VR cues. Kologorov-Smirnov test LEC: VR Off - *n* = 1603 spatial ROIs, VR On - *n* = 3319 spatial ROIs, *D* = 0.1004, *p* = 6.975*e* – 10, MEC: VR Off - *n* = 624 spatial ROIs, VR On - *n* = 2782 spatial ROIs, *D* = 0.2114, *p* = 3.324*e* – 20. *Figure inset*. Mean by mouse. Two-sided paired t-test LEC: *n* = 6 mice, *t* = 3.0285, *p* = 0.0291, MEC: *n* = 5 mice, *t* = 4.7513, *p* = 0.0090. During the VR Off condition the LEC axons had higher specificity than the MEC, Welch’s unequal variances t-test, VR Off: *t* = 3.1158, *p* = 0.0125, but there was no significant difference for the VR On condition: *t* = 1.3347, *p* = 0.2375. Two-factor ANOVA, effect of VR condition (On vs Off), *F* = 10.2643, *p* = 0.0049, axon type *F* = 8.6396, *p* = 0.0088, interaction of type and condition, *F* = 1.2430, *p* = 0.2796. (**F**) Width of the spatially tuned regions represented by the axons was similar, regardless of the VR condition. Kologorov-Smirnov test, LEC: VR Off - *n* = 1962 spatial fields, VR On - *n* = 4244 spatial fields, *D* = 0.0300, *p* = 1.0000, MEC: VR Off - *n* = 791 spatial fields, VR On - *n* = 3866 spatial fields, *D* = 0.0916, *p* = 3.243*e* – 5. *Figure inset*. Mean by mouse. Two-sided paired t-test LEC: *n* = 6 mice, *t* = 0.6121, *p* = 0.5672, MEC: *n* = 5 mice, *t* = 2.3255, *p* = 0.0807. There was no significant difference in place field width between the the LEC and MEC axons Welch’s unequal variances t-test, VR Off: *t* = 1.6951, *p* = 0.1385, VR On: *t* = 0.3193, *p* = 0.7571. Two-factor ANOVA, effect of VR condition (On vs Off), *F* = 3.5167, *p* = 0.0771, axon type *F* = 1.6563, *p* = 0.2144, interaction of type and condition, *F* = 0.6997, *p* = 0.4139. (**G**) *Bottom*. Imaging frame stability for both the LEC and MEC datasets, measured as the mean correlation to the time average was similar for both the LEC and MEC datasets. Welch’s unequal variances t-test *t* = 0.1749, *p* = 0.8679. *Top*. There was no relationship between the percentage of spatially tuned axons and the mean frame stability. Linear regression, LEC: *n* = 6 mice, *r*^2^ = 0.0181, *p* = 0.8294, MEC: *n* = 5 mice, *r*^2^ = 0.0069, *p* = 0.8759. (**H**) The percentage of spatially tuned axons with place field within 50 cm of the goal location is significantly correlated with the mean lick distance to the reward, by mouse for the LEC axons; Linear regression *n* = 6 mice, *r*^2^ = 0.8781, Wald chi-squared test *p* = 0.0060, but not the MEC axons; Linear regression *n* = 5 mice, *r*^2^ = 0.0004, Wald chi-squared test *p* = 0.9749. (**I**) Jaccard overlap score for axons with a spatial field in the reward zone and area just prior (within 50 cm) between VR Off and On contexts. The reward zone was more similarly represented by the LEC axons than the MEC axons, Welch’s unequal variances t-test, LEC: *n* = 6 mice, MEC: *n* = 5 mice, *t* = 6.2490, *p* = 0.0004.

**Figure S4.**
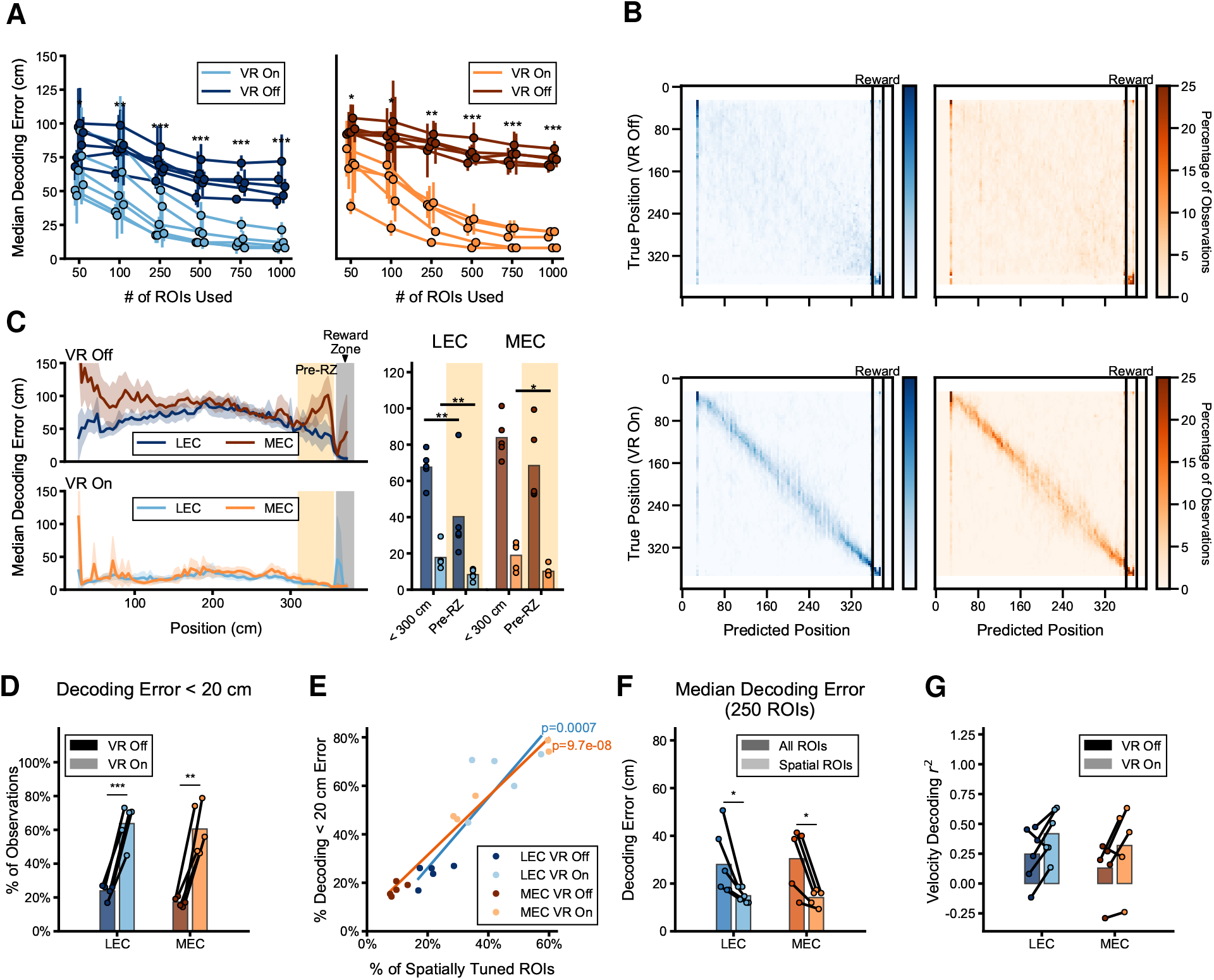
Details on decoding position and velocity. (**A**) Position decoding improved with the addition of more ROIs. *Left*: LEC, *Right*: MEC. Decoding on 3 different random samples of ROIs per animal. Mean value for median error by mouse and 95% confidence interval (bars) shown. Annotations indicate significance between VR On and VR Off conditions. Two-sided paired t-test between VR Off and VR On condition, *n* = 15 LEC ROI sets, *n* = 15 MEC ROI sets: * - *p* < 0.05, ** - *p* < 0.01, *** - *p* < 0.001. (**B**) Confusion matrices for position decoding; successful position decoding heavily relied on visual cues. *Left*: LEC, *Right*: MEC, Top: VR Off, *Bottom*: VR On. (**C**) Position decoding error by position bin was lower at locations near to the reward zone. *Left*: Error by position bin, median decoding error averaged by mouse and randomized ROI sub-sample. Shaded error represents 95% confidence interval. *Right*: Comparison of decoding error far from the reward zone with errors near to the reward zone (within 50 cm). Decoding was significantly improved at locations near to the reward for LEC axons in both VR conditions and for the VR On condition for the MEC axons. Two-sided paired t-test between mouse median error near and far from RZ, VR On, LEC: *n* = 5 mice, *t* = 4.9934, *p* = 0.0075, MEC: *n* = 5 mice, *t* = 3.9685, *p* = 0.0166, VR Off, LEC: *t* = 4.2869, *p* = 0.0128, MEC: *t* = 2.7244, *p* = 0.0527. (**D**) Position decoding success rate as percentage of samples with an error under 20 cm. Bars show all observations, points are mean by mouse. Higher percentage of successful decoding when the VR cues are tuned on. Two-sided paired t-test LEC: *n* = 5 mice, *t* = 9.6725, *p* = 0.006, MEC: *n* = 5 mice, *t* = 6.0122, *p* = 0.0039. (**E**) There was a correlation between the percentage of spatially tuned axons in a field of view and the position decoding results for both the LEC and MEC axon datasets. Linear regression LEC: *n* =10 mouse x VR condition, *r*^2^ = 0.7724, MEC: *n* =10 mouse x VR condition, *r*^2^ = 0.9757, Wald chi-squared test LEC: *p* = 0.0008, MEC: *p* = 9.691*e* – 8. (**F**) Position decoding using 250 of the spatially tuned axons was more successful than decoding position from 250 randomly selected axons. Two sided paired t-test LEC: *n* = 6 mice, *t* = 2.9522, *p* = 0.318, MEC: *n* = 5 mice *t* = 3.5759, *p* = 0.0233. (**G**) Decoding the current velocity was significantly better than a mean velocity model for the LEC axonal projection. Mouse mean *r*^2^, *n* = 6 mice, VR Off: *r*^2^ = 0.2471, VR On: *r*^2^ = 0.4167, one-sided one-sample t-test VR Off: *p* = 0.0236, VR On: *p* = 0.0190 and shows a positive trend in the MEC projection. Mouse mean *r*^2^, *n* = 6 mice, VR Off: *r*^2^ = 0.1298, VR On *r*^2^ = 0.3188, one sided, one sample t-test VR Off: *p* = 0.1486, VR On: *p* = 0.0548. During the VR On condition, there was a non-significant trend for better velocity decoding, especially in the LEC projection. Two-sided paired t-test, LEC: *t* = 2.5466, *p* = 0.0514, MEC: *t* = 2.1220, *p* = 0.1010.

**Figure S5.**
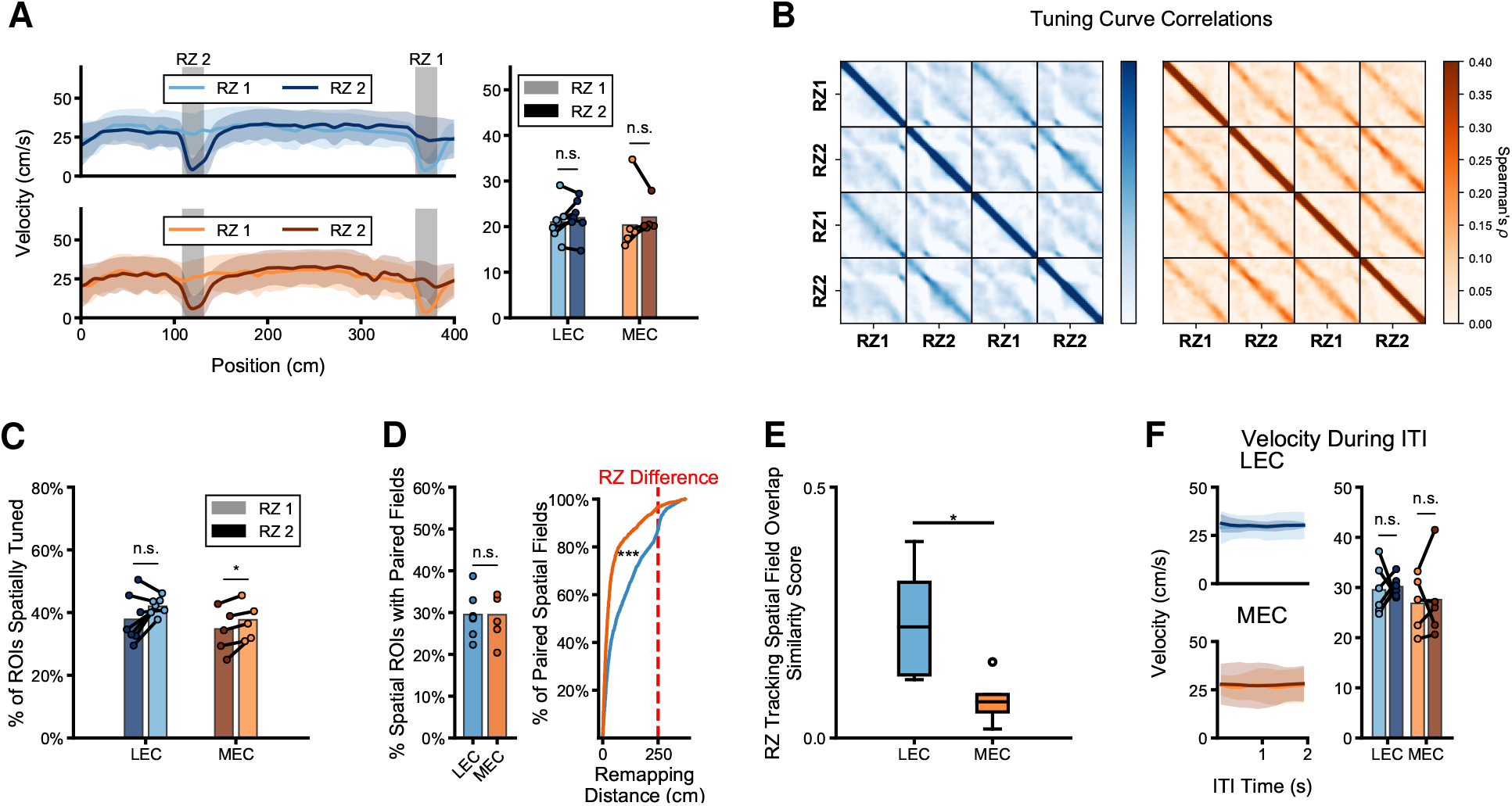
LEC axons activity reflects a change in goal location. (**A**) Velocity during the alternating rewards task. *Left*. Mean velocity at each position for both the RZ1 and RZ2 laps, shaded region depicts standard deviation across mice. *Right*. The mean velocity across all positions for both RZ1 and RZ2 laps was similar. Bars show mean of all observations, points show mean by mouse. Two sided paired t-test LEC: *n* = 7 mice, *t* = 1.0787, *p* = 0.3222, MEC: *n* = 5 mice, *t* = 0.2170, *p* = 0.8388. (**B**) Correlation (Spearman’s *ρ*) between population vectors of mean Ca^2+^ event rate for every position for each reward zone block of the experiment (*Left*. LEC, *Right*. MEC). MEC tuning is more similar regardless of current reward location, while similarity between peri-reward zone representations can be seen in the LEC axonal activity. Mean across mice, LEC: *n* =7 mice, MEC *n* = 5 mice. (**C**) Both LEC and MEC labeled datasets had a similar number of spatially tuned axons in both RZ1 and RZ2 conditions, with slightly more spatially tuned ROIs in the MEC dataset during the RZ2 condition. Bars represent percent of all ROIs LEC: 37.8% RZ1, 42.0% RZ2, MEC: 34.8% RZ1,37.7% RZ2. Points show individual mice, Two sided paired t-test, LEC: *n* = 7 mice, *t* = 1.8013, *p* = 0.1217, MEC: *n* = 5 mice, *t* = 2.9350, *p* = 0.0426. (**D**) LEC tuned axons tended to remap further following a change in reward location than the tuned MEC axons. *Left*. The percentage of spatially tuned ROIs that were tuned in both the RZ1 and RZ2 conditions. Not significantly different between LEC and MEC datasets, LEC: 29.5%, MEC: 29.5%, Welch’s unequal variances t-test, *n* = 7 LEC, *n* = 5 MEC, *t* = 0.2935, *p* = 0.7762. *Right*. The difference in centroid location during RZ1 and RZ2 laps, for all ROIs that were spatially tuned in both conditions. LEC spatially tuned axons moved much further than MEC, Kologorov-Smirnov test *n* = 1601 LEC ROIs, *n* = 1236 MEC ROIs, *D* = 0.2870, *p* = 5.551*e* – 16, red line indicates distance between reward locations. (**E**) The reward zone was more similarly represented by the LEC axons than the MEC axons. Jaccard overlap score for ROIs tuned within the reward location and 50 cm prior, LEC: *n* = 7 mice, mean overlap score = 0.2214, MEC: *n* = 5 mice, mean overlap score = 0.0721, Welch’s unequal variance t-test *t* = 3.0719, *p* = 0.0137. (**F**) Mice maintained similar velocities during the ITI, regardless of the current reward location. RZ1 shown in the lighter color, RZ2 the darker *Left, Top*. Mean velocity (line) and 95% confidence interval (shaded) for LEC animals, *Left, Bottom*. for MEC. *Right*. Mean velocity for the entire ITI, bars represent all data, points show the mean by mouse, LEC: 29.5 cm/s RZ1,30.2 cm/s RZ2, *n* =6 mice, two sided paired t-test, *t* = 0.2555, *p* = 0.8085, MEC: 26.8 cm/s RZ1,27.6 cm/s RZ2, *n* = 5 mice, two sided paired t-test, *t* = 0.2638, *p* = 0.8049.

**Figure S6.**
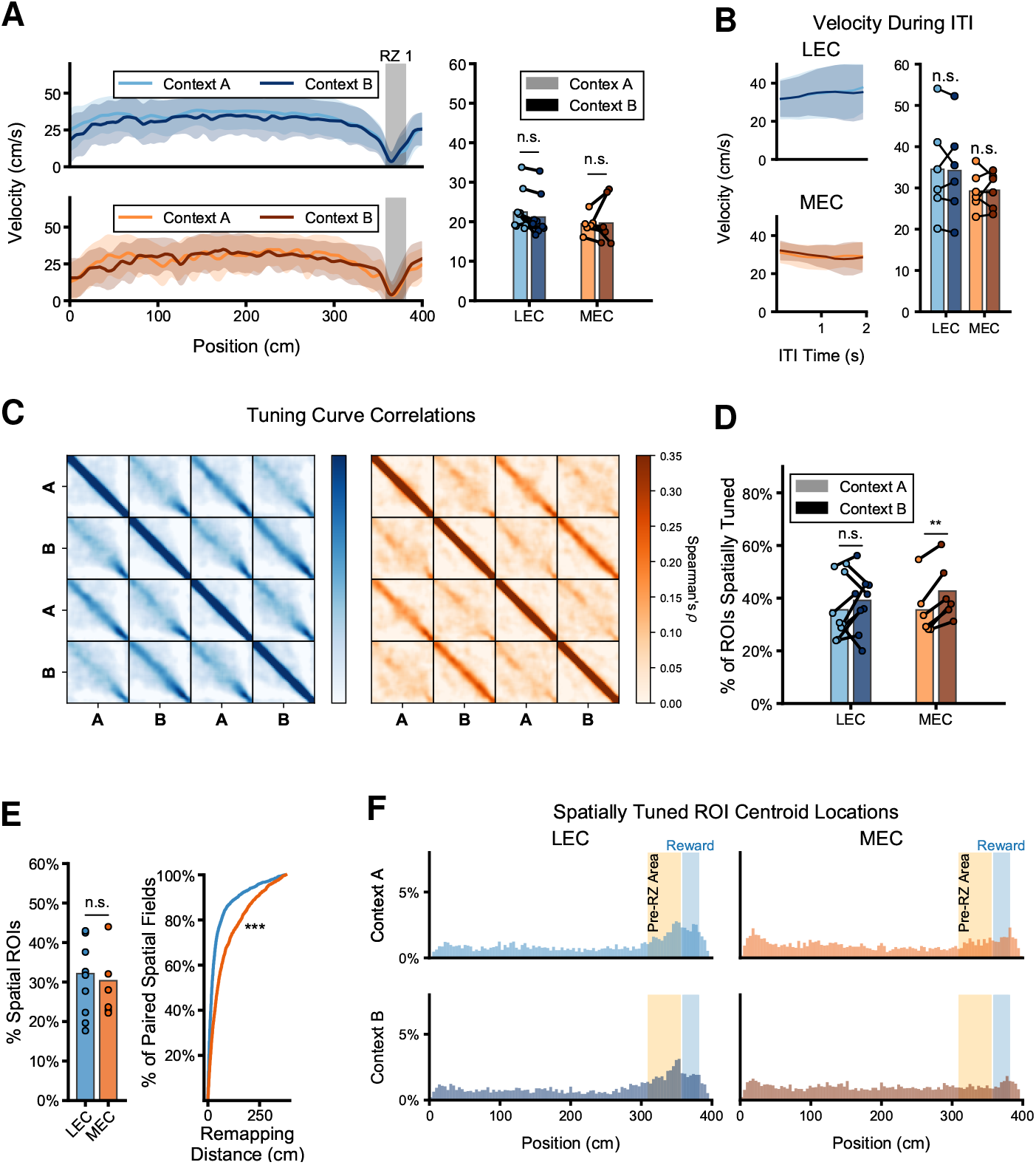
MEC and LEC axon activity shift differently during context alternation task. (**A**) Velocity during the context change task. *Left*. Mean velocity at each position for both the Context A and Context B laps, shaded region depicts standard deviation across mice. *Right*. The mean velocity across all positions for both Context A and Context B laps was similar. Bars show mean of all observations, points show mean by mouse. Two-sided paired t-test LEC: *n* = 9 mice, *t* = 1.8010, *p* = 0.1094, MEC: *n* = 6 mice, *t* = 0.3690, *p* = 0.7272. (**B**) Mice maintained similar velocities during the ITI, regardless of the Context they just experienced. Context A shown in the lighter color, Context B the darker *Left, Top*. Mean velocity (line) and 95% confidence interval (shaded) for LEC animals, *Left, Bottom*. for MEC. *Right*. Mean velocity for the entire ITI, bars represent all data, points show the mean by mouse, LEC: 34.5 cm/s Context A, 34.2 cm/s Context B, *n* =6 mice, two-sided paired t-test, *t* = 0.1955, *p* = 0.8527, MEC: 29.2 cm/s Context A, 29.5 cm/s Context B, *n* = 6 mice, two-sided paired t-test, *t* = 0.1465, *p* = 0.8892. (**C**) Correlation (Spearman’s *ρ*) between population vectors of mean Ca^2+^ event rate for every position for each RZ block of the experiment (Left. LEC, *Right*. MEC). LEC tuning was more similar at most location between contexts, but especially near the RZ. Mean across mice, LEC: *n* = 9 mice, MEC *n* = 6 mice. (**D**) Both LEC and MEC had similar numbers of spatially tuned axons in both Context A and Context B conditions, with a slightly more spatially tuned ROIs in the MEC dataset during Context B. Bars represent percent of all ROIs LEC: 35.6% Context A, 39.2% Context B, MEC: 35.5% RZ1,42.7% RZ2. Points show individual mice, Two-sided paired t-test, LEC: *n* = 9 mice, *t* = 0.727, *p* = 0.5022, MEC: *n* = 6 mice, *t* = 5.6667, *p* = 0.0024. (**E**) Spatially tuned MEC axons tended to remap further between contexts than LEC axons. *Left*. The percentage of spatially tuned ROIs that were tuned in both the Context A and B. Not significantly different between LEC and MEC datasets, LEC: 29.5%, MEC: 29.5%, Welch’s unequal variances t-test, LEC: *n* = 9 mice, MEC: *n* =6 mice, *t* = 0.3746, *p* = 0.7146. *Right*. The difference in centroid location between Context A and B laps, for all ROIs that were spatially tuned in both conditions. LEC spatially tuned axons moved much further than MEC, Kologorov-Smirnov test *n* = 1898 LEC ROIs, *n* = 1621 MEC ROIs, *D* = 0.2635, *p* = 3.449*e* – 53. (**F**) Percentage of all spatially tuned fields with a centroid at each location. An enrichment of LEC axons with spatial fields in the 50 cm area prior to the start of the reward zone (highlighted). *Top*. Context A *Bottom*. Context B.

**Figure S7.**
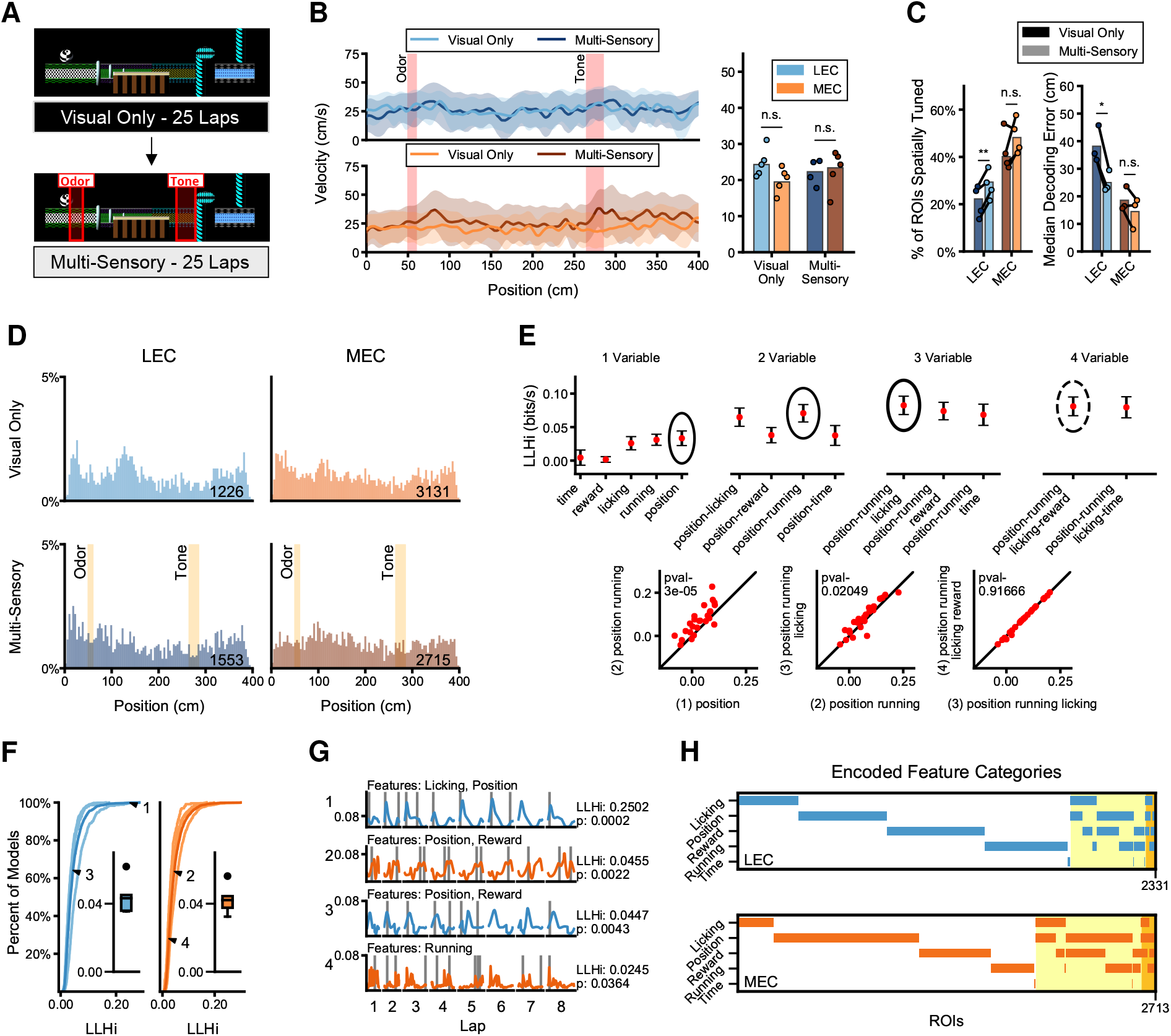
EC Axons responses when reward location is different on every lap. (**A**) Diagram of the multi-sensory variant of the shuffled rewards task. Spatially fixed odor (carvone) was present at 50 - 60 cm and tone (5 kHz) was placed at 265 - 285 cm. (**B**) Velocity during the random foraging task was similar between LEC and MEC labeled mice. *Left*. Mean velocity at each position, shaded region depicts standard deviation between mice. *Right*. Quantification, by mouse of the mean velocity for the entire track was not significantly different between the LEC and MEC mice, Welch’s unequal variances t-test LEC: *n* = 5 mice, MEC: *n* = 5 mice, visual cues only *t* = 2.0237, *p* = 0.0782, multi-sensory task *t* = 0.1789, *p* = 0.8632. (**C**) *Left*. Slightly more axons were determined to be significantly spatially tuned (*p* < 0.05) during laps when there were spatially fixed salient tone and odor cues LEC: 22.2%, Visual Only, 29.3%, *n* = 4, Multi-Sensory, two-sided paired t-test *t* = 5.9452, *p* = 0.0095, MEC: 40.1%, Visual Only, 48.2%, Multi-Sensory *n* = 4 mice, *t* = 1.6976, *p* = 0.1882. *Right*. Position decoding error was also lower when spatially fixed salient cues were present. Median decoding error averaged by mouse LEC: 33.7 cm Visual Only, 21.8 cm Multi-Sensory, *n* = 3 mice, two sample paired t-test *t* = 5.8919, *p* = 0.0276, MEC: 16.4 cm Visual Only, 12.0 cm Multi-Sensory *n* = 3 mice, two sample paired t-test *t* = 10.0000, *p* = 0.0099. (**D**) Percentage of all spatially tuned fields with a centroid at each location during random foraging task. *Top*. First 25 laps with only the visual VR cues, LEC: *n* = 5 mice, MEC: *n* = 5 mice. *Bottom*. Spatially spatial fields during the following 25 laps when spatially fixed odor and tone cues were present, LEC: *n* = 4 mice, MEC: *n* = 4 mice. (**E**) *Top*. Example of model selection process for a single axon (mean ± SEM LLHi in bits/second). Best fit models are circled. Bold circle represents the simplest model which significantly (*p* < 0.05) outperforms any simpler model. *Bottom*. Procedure for identifying model significance using best fits from the *top* panels. (**F**) LLHi over mean firing rate for all best fit models determined by the model selection process (only significant models *p* < 0.05 included). Thin lines - individual mice, Thick lines - all axons *Left*. LEC. *Right*. MEC. *Inset*. LLHi Average by mouse. (**G**) Example of best-fit models for 4 axons during 10 laps with a verity of model fit successes. Colored lines show the predicted event probability and gray lines show recorded Ca^2+^ event times. Axes labels 1 - 4 correspond to annotations in **G**. (**H**) Features categories used in the Best fit models of for each of the axons recorded during the random foraging task. Highlighted portion shows mixed-selective models.

**Figure S8.**
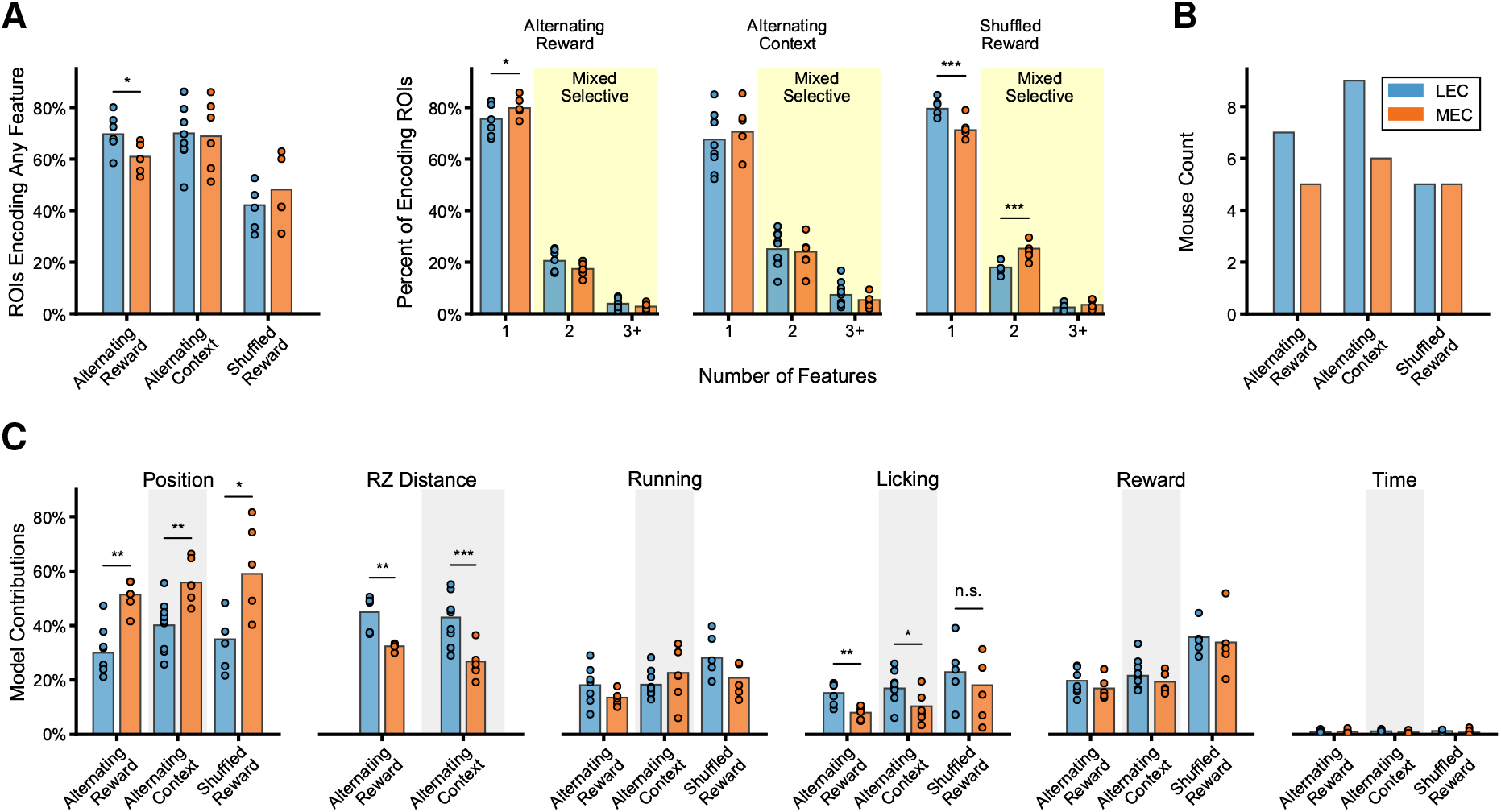
Distance to reward is a better at predicting more activity in more LEC axons, while context/position is more predictive of MEC axonal activity. (**A**) *Left*. Comparison of logistic regression models across different tasks. Successful models show a significant (*p* < 0.5) improvement over a mean firing rate model across all laps. Similar percentages of LEC and MEC axons are successfully modeled for each of the tasks, Welch’s unequal variances t-test. *Right*. Percentages of best-fit models incorporating a single type of features or multiple (mixed selected), significance shown is interaction between axon type and number of features encoded, linear mixed effects model with axons type and number of features as fixed factors and mouse as the random factor. * - *p* < 0.05, *** - *p* < 0.001 (**B**) Number of mice of each axon type used for logistic regression analysis. Due to either uncorrectable imaging or not running sufficient laps, there were different numbers of mice for each of the experiments. (**C**) Comparison of logistic regression models between tasks. Selected predictors influencing models for each task; context-position was consistently influential for more of the MEC axons, while distance to reward significantly influenced a higher percentages of LEC models. Welch’s unequal variances t-test between subregion for percentage of best fit models including each feature (by mouse) for each task. * - *p* < 0.05, ** - *p* < 0.01, *** *p* < 0.001.

**Figure S9.**
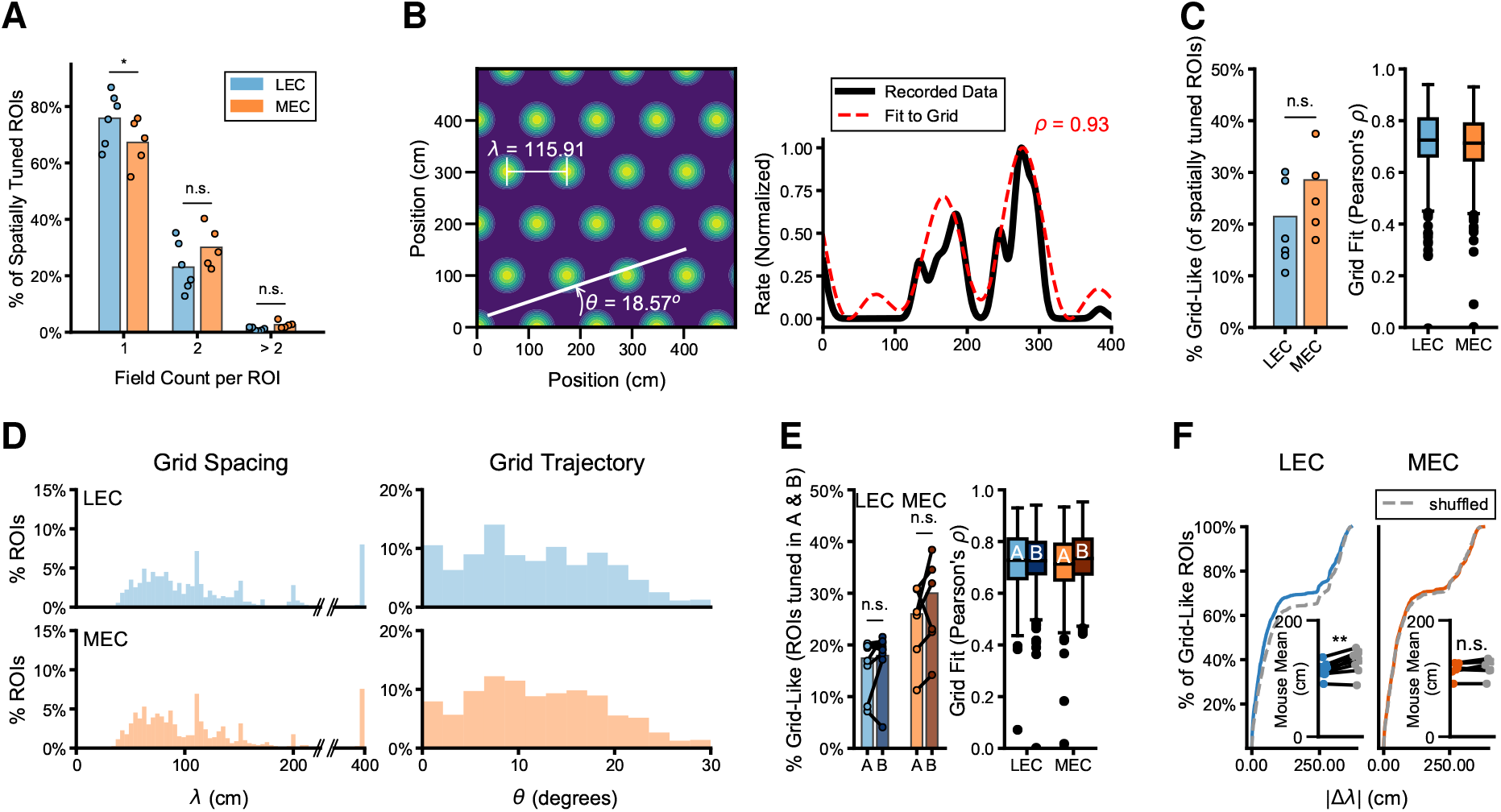
Fitting EC responses as a 1D trajectory though a 2D grid. (**A**) Spatially tuned MEC axons tended to have increased field counts during VR navigation as compared to LEC. Mean field count LEC: 1.2609 ± 0.0130, *n* = 4922 tuned axons, MEC: *n* = 3406 tuned axons, 1.3673 ± 0.0181. Linear mixed effects model with axon type and spatial field count as fixed factors and mouse as the random factor, *n* = 6 mice LEC, *n* = 5 mice MEC, axon type x single place field interaction, *z* = 2.011, *p* = 0.0443, axon type x two place fields *z* = 1.645, *p* = 0.1000, axon type x > two place fields *z* = 0.366, *p* = 0.7142. (**B**) Some axons have tuning curves which are fit well to models of a 1D slice through a 2D grid tuning field. *Left*. 2-dimensional grid that matches the tuning curve for an example MEC axon recording. *Right*. Tuning curve for the example grid-like MEC axon shown to the left. Recorded tuning curve in black, idealized grid-fit in red.(**C**) *Left*. There were not significantly more MEC axons classified as “grid-like” compared to the LEC (VR On condition). LEC: 21.4%, and MEC: 28.5%, Welch’s unequal variance t-test by mouse, LEC: *n* = 6 mice, MEC: *n* = 5 mice, *t* = 1.3347, *p* = 0.2160. *Right*. There was also no evidence that the “grid-like” activity recorded from MEC axons was more likely to be from a 1D trajectory through a 2D grid cell by comparing fits to ideal grid-derived tuning curves, by mouse mean observed tuning curve correlation to ideal, LEC: *n* =6 mice, MEC: *n* = 5 mice, Welch’s unequal variances *t* = 1.5273, *p* = 0.1610. (**D**) Best fit values for the ideal grid spacing (λ, *left*) and trajectory (*θ*, *right*) for EC axon tuning curves classified as “grid-like”. (**E**) *Left*. Percentage of both LEC and MEC axons classified as “grid-like” for both contexts. LEC: 17.4% Context A, 17.9% Context B, MEC: 26.0% Context A, 30.0% Context B. Between context comparison, two sample paired t-test, LEC: *n* = 9 mice, *t* = 1.1528, *p* = 0.2823, MEC: *n* = 6 mice, *t* = 1.3022, *p* = 0.2496. Significantly more “grid-like” axons were observed in the MEC axons in Context B as compared to LEC, but not in Context A. Welch’s unequal variances t-test of mouse mean, Context A: *t* = 2.1495, *p* = 0.0633, Context B: *t* = 2.3585, *p* = 0.0490. *Right*. We cannot resolve that the “grid-like”activity recorded from MEC axons was a better fit for a 1D trajectory through a 2D grid than the LEC axons. Welch’s unequal variances t-test *n* = 9 mice LEC, *n* = 6 mice MEC, Context A: *t* = 1.6248, *p* = 0.1047, Context B: *t* = 1.0461, *p* = 0.2959. (**F**) The difference in the calculated grid spacing, λ, for all axons classified as potentially “grid-like” compared to shuffled values. *δ*λ was significantly lower for LEC axons, but not for MEC axons indicating no evidence for a grid cell biased projection, LEC: *n* = 1768, *D* = 0.0854, *p* = 4.953*e* – 6, MEC: *n* = 1413, *D* = 0.0234, *p* = 0.83358. *Inset*. Mean by mouse, compared to shuffled data, two-sided, two sample t-test LEC: *n* = 9 mice, *t* = 4.6151, *p* = 0.0017, MEC: *n* = 6 mice, *t* = 1.1432, *p* = 0.3047.

### Description of fitting EC axon activity observed in CA1 to grid-cell model during Virtual Navigation and Context Change tasks

During the VR navigation task, we did observe a tendency for the spatially tuned MEC axons to have more place fields per ROI (Figure S9A). This led us to check the activity of the axons for hallmarks of grid-cell activity. Using the LEC projection as a control, since LEC axons might have spatially tuned fields but not grid cell like activity, we found no significant evidence for grid tuning in the MEC projection to CA1. Specifically, applying a previously established classification scheme to search for grid-like tuning on a linear track (Yoon et al. 2016, Gu et al. 2018, Cholvin et al. 2021), we found no significant difference in the proportion of MEC axons showing periodic spatial tuning compared to LEC (LEC 21.45%, MEC 28.52%, *p* = 0.2160 unequal variances t-test, Figure S9C). We then fit all of the grid-like ROIs to the 2D grid cell lattice to see how well the spacing of potential grid fields matches expected grid cell firing rates. Notably, the fits for recordings of MEC axons were no better than for LEC axons (Person’s *ρ*, LEC: 0.7250, MEC: 0.7131, Figure S9B, C). Further, we did not observe any greater evidence for clustering in the grid-spacing (λ) or grid-angle (*θ*) in the MEC as compared to LEC axons–which, if present, could have indicated activity consistent with input from individual grid modules (Figure S9D). We conclude, in agreement with previous reports (Cholvin et al. 2021), that while we cannot rule out the possibility of grid tuning in the MEC-to-CA1 projection, the majority of spatially tuned axons projecting from the EC to CA1 are likely to be spatial/non-grid cells (just as the majority of tuned cells in the MEC are Diehl et al. (2017)).

In the alternating context task, we observed a higher percentage of “grid-like” spatially tuned axons in the MEC compared to the LEC projection; nevertheless, the population of LEC ROIs which also passed the “grid-like” classification criteria was larger than expected and likely indicates false-positive detections (LEC, Context A: 17.4%, Context B: 17.9%, MEC Context A: 26.0%, Context B: 30.0%, Figure S9E). In order to check how well the “grid-like” axons matched grid tuning, we fit both the LEC and MEC axons’ tuning curves to 1D slices of idealized grid fields. We found no difference in how well they fit, on average (mean grid fit, *ρ*, LEC Context A: 0.7275, Context B: 0.7256, MEC Context A: 0.7131, Context B: 0.7345, Figure S9E), suggesting that the grid-like activity might be due to the periodic nature of the virtual cues (some are repeated along the track) and the fixed reward zone. Additionally, grid cells would be expected to have the same grid field spacing (λ) regardless of the context, so we compared how much the grid spacing changed between contexts. We found that while the difference in the LEC grid-fit spacing was altered slightly less than would be expected by chance (likely due to the bias of LEC ROIs to form spatial fields around the reward zone), the MEC grid fits were identical to the shuffled data (mean by mouse, *δλ* compared to shuffled λ all ROIs, LEC *p* = 0.0017, MEC *p* = 0.3047, Figure S9F). These results are in agreement with what we report from the single VR context task, in that we see no specific evidence that the majority of MEC axons projecting to CA1 come from grid-cells.

